# FRUITFULL-like genes regulate flowering time and inflorescence architecture in tomato

**DOI:** 10.1101/2020.09.28.316562

**Authors:** Xiaobing Jiang, Greice Lubini, José Hernandes-Lopes, Kim Rijnsburger, Vera Veltkamp, Ruud A. de Maagd, Gerco C. Angenent, Marian Bemer

**Author notes:** The author responsible for distribution of materials integral to the findings presented in this article in accordance with the policy described in the Instructions for Authors (www.plantcell.org) is: Marian Bemer.

## Abstract

The timing of flowering and inflorescence architecture are critical for the reproductive success of tomato, but the gene regulatory networks underlying these traits have not been fully explored. Here we show that the tomato *FRUITFULL*-like (*FUL*-like) genes *FUL2* and *MADS-BOX PROTEIN 20 (MBP20)* induce flowering and repress inflorescence branching by promoting floral meristem determinacy. *FUL1* fulfils a less prominent role and appears to depend on FUL2 and MBP20 for its upregulation in the inflorescence- and floral meristems. *MBP10*, the fourth tomato *FUL*-like gene, has probably lost its function. The tomato FUL-like proteins cannot homodimerize in *in vitro* assays, but heterodimerize with various other MADS-domain proteins, potentially forming distinct complexes in the transition meristem and floral meristem. Transcriptome analysis of the primary shoot meristems revealed various interesting downstream targets, including four repressors of cytokinin signalling that are upregulated during the vegetative-to-reproductive transition in *ful1 ful2 mbp10 mbp20* mutants. FUL2 and MBP20 can also bind *in vitro* to the upstream regions of these genes, thereby probably directly stimulating cell division in the meristem upon the transition to flowering. The control of inflorescence branching does not occur via the CKXs, but appears regulated by repression of transcription factors such as *TM3, APETALA 2b* (*AP2b*) and *AP2a*.

## INTRODUCTION

The MADS-box transcription factor gene family is involved in almost every developmental process in plants (Smaczniak et al., 2012a), and the members of the angiosperm-specific APETALA1/FRUITFULL (AP1/FUL) subfamily play key roles in flowering and fruit development (Litt and Irish, 2003; McCarthy et al., 2015). In the core eudicots, the AP1/FUL subfamily consists of three clades, euAP1, euFULI and euFULII (Litt and Irish, 2003). The Arabidopsis genome carries four *AP1/FUL*-clade genes, with *AP1* functioning as an A-class gene in the ABC model, promoting perianth identity (Mandel et al., 1992; Theissen and Saedler, 2001) and establishing floral meristem identity and floral initiation (Ferrándiz et al., 2000; Kaufmann et al., 2010). Its lower-expressed paralog CAULIFLOWER (*CAL*) functions to a large extent redundantly with *AP1* (Bowman et al., 1993; Ye et al., 2016). The euFULII-clade gene *AGL79* appears to have a minor function in roots (Gao et al., 2018), whereas the euFULI gene *FUL* is a pleiotropic gene with important roles in fruit development and flowering. *FUL* regulates many aspects of flowering in Arabidopsis, including flowering time (Ferrándiz et al., 2000; Melzer et al., 2008), repression of inflorescence meristem (IM) identity (together with *AP1/CAL*; Ferrándiz et al., 2000), inflorescence architecture (Bemer et al., 2017), axillary inflorescence outgrowth (together with *SOC1*; Karimi et al., 2020) and IM termination (Balanzà et al., 2018). In a wide range of angiosperm species, *FUL*-like genes regulate fruit development (Gu et al., 1998; Jaakola et al., 2010; Bemer et al., 2012; Pabón-Mora et al., 2012; Zhao et al., 2019) and flowering (Ferrándiz et al., 2000; Berbel et al., 2012; Pabón-Mora et al., 2013; Ping et al., 2014; Jaudal et al., 2015; Jia et al., 2015; Li et al., 2019; Zhang et al., 2021). In the model crop species tomato (*Solanum lycopersicum*), the euFULI-clade genes *Solanum lycopersicum FUL1* (*SlFUL1/TM4/TDR4*, hereafter called *FUL1*) and *SlFUL2* (*FUL2/MBP7*, hereafter called *FUL2*) play important roles in fruit development and ripening (Bemer et al., 2012; Shima et al., 2013; Wang et al., 2019), but flowering phenotypes have not yet been described for these genes, nor for the tomato euFULII-clade genes MADS-BOX PROTEIN 10 (MBP10) and MBP20. This is remarkable given the strong upregulation of *FUL1, FUL2* and *MBP20* expression during the transition from shoot apical meristem (SAM) to inflorescence- and floral meristem (Park et al., 2012).

Flowering is an important agricultural trait in tomato, as both the onset and termination of flowering, as well as inflorescence architecture, determine crop yield. Tomato is also an interesting model species considering its sympodial inflorescence architecture, which is distinct from that of the monopodial Arabidopsis inflorescence. While the Arabidopsis shoot apical meristem (SAM) develops into an inflorescence meristem (IM), which subsequently forms floral meristems (FM) on its flank (indeterminate inflorescence), the tomato SAM is doming to form the transition meristem (TM) that terminates directly into an FM (determinate inflorescence), but forms a new IM on its flank. This iterative process results in a zigzagged inflorescence (Lippman et al., 2008). Moreover, tomato has a compound shoot, which resumes vegetative growth from the axillary meristem of the youngest leaf when the first IM terminates. After activation of the axillary meristem (then called sympodial meristem, SYM), the shoot forms three leaves before terminating again into the first flower of the second inflorescence (Pnueli et al., 1998; Szymkowiak and Irish, 2006; Lippman et al., 2008), upon which a new axillary meristem takes over vegetative growth. This process forms the compound shoot, where three leaves and an inflorescence comprise a sympodial unit, a pattern that is endlessly repeated in the wild-type tomato.

The genes that regulate these flowering processes have been very well studied in the indeterminate Arabidopsis inflorescence, where the floral integrators FLOWERING LOCUS T (FT) and SUPPRESSOR OF OVEREXPRESSION OF CONSTANS1 (SOC1) induce the transition to flowering, after which *TERMINAL FLOWER* 1 (*TFL1*) determines IM fate by repressing FM genes such as *LEAFY* (*LFY*) and *AP1* (Sablowski, 2007; Lee and Lee, 2010; Serrano-Mislata et al., 2017; Zhu et al., 2020), while AP1 on its turn represses *TFL1*, so that clear borders between the FM and IM meristems are achieved (Liu et al., 2013; Goslin et al., 2017). The variation in inflorescence structures of different species can be largely explained by different temporal and spatial expression of flower-repressing *TFL* homologs and flower-inducing *LFY/AP1/FUL* homologs (McGarry and Ayre, 2012; Périlleux et al., 2019). In legumes for example, the indeterminate inflorescence does not form FMs on its flank, but secondary IMs, due to repression of the *TFL*-homolog in these meristems by the euFULII clade proteins VEG1 (pea) or MtFUL1-c (Medicago) (Berbel et al., 2012; Cheng et al., 2018; Zhang et al., 2021). These secondary IMs do form FMs, resulting in a compound inflorescence (Benlloch et al., 2015).

In tomato, *SINGLE FLOWER TRUSS* (*SFT*), *FALSIFLORA* (*FA*) and *MC* are essential for the transition to flowering and control of floral meristem identity similar to their orthologs *FT, LFY* and *AP1* in Arabidopsis (Molinero□Rosales et al., 1999; Molinero-Rosales et al., 2004; Yuste-Lisbona et al., 2016). However, mutants in the *TFL*-ortholog *SELF PRUNING* (*SP*), which is not expressed in the primary SAM/VM, but highly expressed in early axillary meristems (Thouet et al., 2008), loose this growth indeterminacy and terminate their SYMs early, resulting in termination of growth after a few sympodial units (Pnueli et al., 1998). Inflorescence architecture is normal in *sp* mutants (Pnueli et al., 1998). Instead, tomato inflorescence architecture is influenced by other factors that regulate the establishment or timing of FM maturation/termination (Lippman et al., 2008). Delayed meristem maturation, occurring in the mutants *compound inflorescences* (*s*), *falsiflora* (*fa*; *FA is the* ortholog of *LFY*) and *anantha* (*an*), induces additional IM formation before the determinate FM has matured (Szymkowiak and Irish, 2006; Chae et al., 2008; Lippman et al., 2008; Souer et al., 2008; Park et al., 2012; Soyk et al., 2017), resulting in branched (compound) inflorescences. Conversely, precocious activation of the AN/FA complex leads to early FM induction and thus early flowering (MacAlister et al., 2012). In addition to these factors, several MADS-domain transcription factors function in tomato flowering, mainly in conferring IM or FM identity. In the *jointless* (*j*) and *macrocalyx* (*mc*) mutants, flowering is delayed and the inflorescence reverts to vegetative growth after a few flowers, probably because the meristems adopt SYM identity instead of FM identity (Szymkowiak and Irish, 2006; Yuste-Lisbona et al., 2016). Mutations in the SEPALLATA-like genes *JOINTLESS 2* (*J2/SlMBP21*), *ENHANCER OF JOINTLESS 2* (*EJ2/MADS1*) and the *SOC1*-like genes *TM3* and *SISTER OF TM3* (*STM3*) affect inflorescence branching through yet uncharacterized mechanisms (Gomez Roldan et al., 2017; Soyk et al., 2017; Soyk et al., 2019; Alonge et al., 2020). Interestingly the *tm3 stm3* mutations suppress the enhanced branching phenotype of the *j2 ej2* mutant (Alonge et al., 2020), suggesting that these MADS-domain TFs have an opposite function in FM development. Natural mutations or structural variants in several of these MADS-box genes have been important for domestication, either through the regulation of pedicel abscission (*MC, J, SlMBP21/J2*) or inflorescence architecture (J2, EJ2, TM3, STM3) (Nakano et al., 2012; Liu et al., 2014; Gomez Roldan et al., 2017; Soyk et al., 2019; Alonge et al., 2020). Thus, although tomato inflorescence development differs fundamentally from that of monopodial species such as Arabidopsis, the orthologs of several important Arabidopsis flowering and inflorescence meristem genes are also essential in tomato. However, it is yet unclear if and how the *FUL*-like genes (*SlFUL*)function in the tomato flowering regulatory network.

Here, we investigated the developmental roles of the four *FUL*-like genes in tomato by CRISPR/Cas9-mutagenesis and transcriptome profiling. We demonstrate that sub-functionalization has occurred after duplication within the Solanaceae euFULI and euFULII clades, and that the *FUL1* sequence has undergone further divergence during tomato domestication and breeding. *FUL2* and *MBP20* are highly expressed in the meristem during the vegetative-to-reproductive transition to additively promote tomato flowering and to repress inflorescence branching together with *FUL1*. Transcriptome analysis in the *ful1 ful2 mbp10 mbp20* quadruple mutant revealed that the *FUL*-like genes act parallel to, or downstream of, previously described key regulators such as SFT, FA and AN during both the VM-to-TM transition and the establishment of inflorescence architecture. Instead, our target gene analysis revealed that the delay in transition to flowering may be explained by reduced cytokinin signalling as result of upregulation of *CKX* genes, while the increased branching is probably caused by delayed FM termination as a result of specific MADS-domain and AP2-like transcription factors that are upregulated in the mutant.

## RESULTS

### Expression patterns and protein-protein interaction profiles differ between the tomato FUL homologs

To investigate to what extent the different *FUL*-like genes may have overlapping functions in specific organs, we performed expression profiling using qRT-PCR in the cultivar Moneyberg (Figure 1A). The results show that *FUL1* and *FUL2* are expressed very weakly during vegetative growth and have increased expression in the inflorescence meristem. Their expression remains high throughout reproductive development, where both genes show considerable expression in all floral whorls and all stages of fruit development. *FUL2*, in particular, is strongly expressed in all floral organs, early fruits and ripening fruits, while *FUL1* expression is moderate until the fruit ripening phase, when it increases strongly as reported previously (Bemer et al., 2012). Our data also show the striking differences in the spatial expression of *MBP10* and *MBP20* that are both much more weakly expressed than *FUL1* and *FUL2* in the reproductive tissues, except for the expression of *MBP20* in the inflorescence meristem. *MBP10* especially was extremely weakly expressed, with detectable levels in stem and flower bud only.

**Figure 1.**
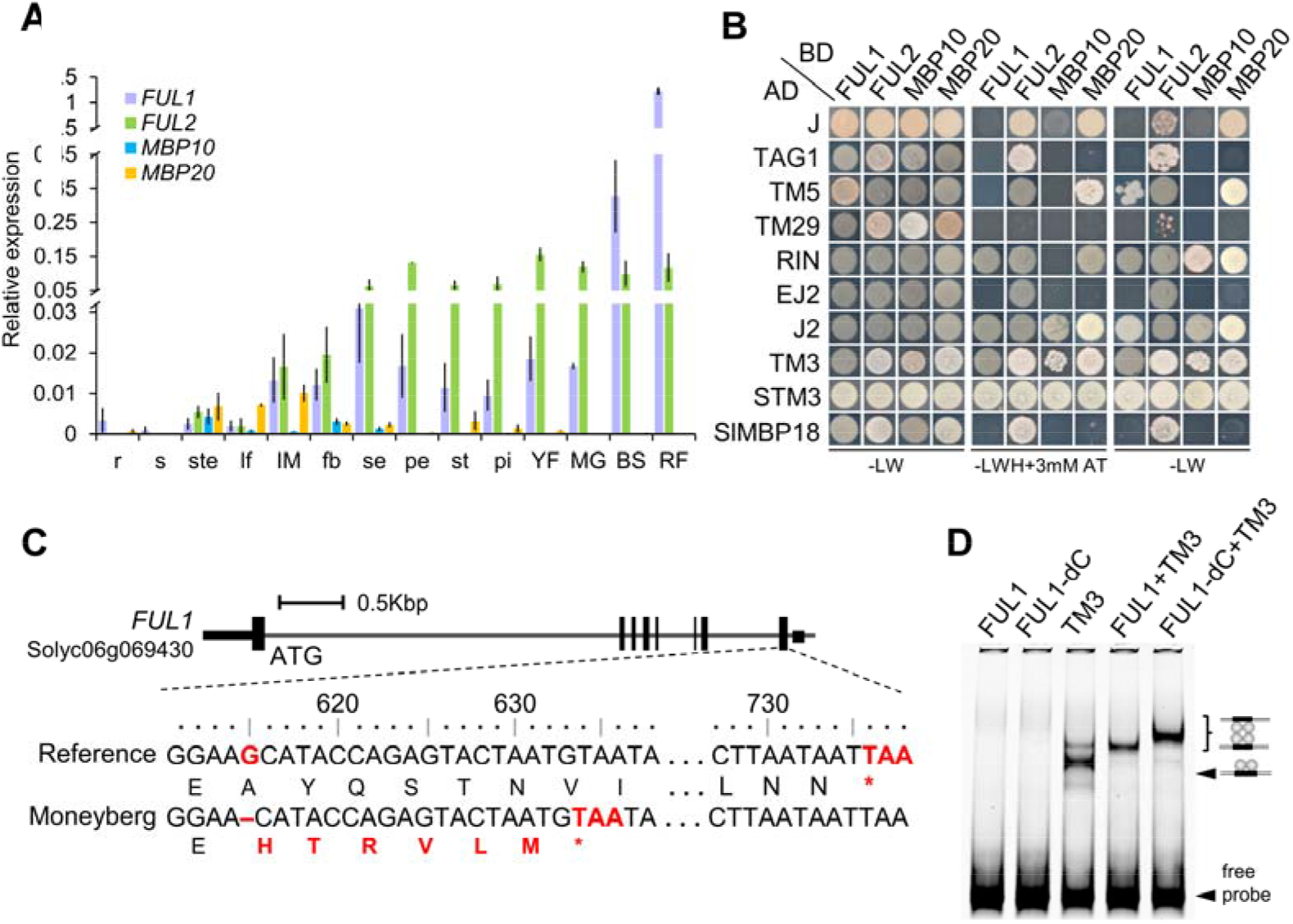
Characterization of the tomato *FUL*-like gene*s*. **(A)** Relative expression profiles of *SlFUL* genes in different organs obtained by qRT-PCR. r: two-week-old root; s: two-week-old shoot; ste: stem below apex; lf: young leaf; IM: inflorescence meristem (dissected apex); fb: closed flower bud; se: sepal; pe: petal; st: stamen; pi: pistil; YF: young fruit; MG: mature green fruit; BR: breaker stage fruit; RR: red ripe fruit. The error bars indicate ±SD based on three biological replicates. **(B)** Yeast two-hybrid assays showing the protein interactions of the FUL-like proteins with other tomato MADS-domain proteins. L, leucine; W, tryptophan; H, histidine; A, adenine; 3-AT, 3-amino-1,2,4-triazole. **(C)** The position of the one-nucleotide deletion present at the 3’ end of the *FUL1* gene in the cultivar Moneyberg, resulting in a protein lacking the C-terminus. The red letters indicate the differences in amino acids, the red nucleotides indicate the stop codons, the vertical black bars are exons. The dotted line refers to what is expanded in the sequence below (part of the last exon). **(D)** EMSA assays showing that the FUL1 truncated protein (FUL1-ΔC) can form higher-order complexes *in vitro*.

MADS-domain transcription factors regulate multiple developmental processes by forming dimeric complexes (De Folter et al., 2005). To investigate which protein complexes may be formed by the different FUL-like proteins, we performed a yeast-two hybrid (Y2H) assay to identify the interactions of FUL1, FUL2, MBP10 and MBP20 with other tomato MADS-domain family proteins (Figure 1B, Supplemental Figure 1). We chose a set of proteins homologous to MADS-domain proteins known to interact with Arabidopsis FUL (De Folter et al., 2005). While Arabidopsis FUL (AtFUL) can form homodimers (Smaczniak et al., 2012b), none of the tomato FUL-like proteins possessed this capacity, nor could they heterodimerize with each other (Supplemental Figure 1). Of the tested MADS-domain proteins, MC, MBP9, MBP12, MBP13, MBP14, MBP22 and MBP24 did not interact with any of the tomato FUL homologs. Our screen showed that the SlFUL proteins could interact with nine other tomato MADS-domain proteins, and we observed clear differences between the interaction profiles of FUL1, FUL2, MBP10 and MBP20. All FUL-like proteins interacted with JOINTLESS (J), J2 (or SlMBP21, hereafter called J2), TM3 and STM3, the co-orthologs of the Arabidopsis flowering integrator SOC1 (Alonge et al., 2020) and the fruit ripening regulator MADS-RIN. FUL2 exhibits the most extensive interaction network, interacting with nine proteins. It is the only protein that strongly interacts with the SEPALLATA (SEP)-like proteins EJ2 and LeSEP1/TM29, and the AGAMOUS (AG) ortholog TAG1 (Figure 1B and Supplemental Figure 1). The latter interaction is especially interesting in the light of the high expression of *FUL2* in pistils, where FUL2 may have a specific function in a complex with the co-expressed TAG1. In addition to these specific interactions, FUL2, FUL1 and MBP20 share interactions with the SEP-like protein TM5 and the AGL42-homolog FOREVER YOUNG FLOWER-LIKE (FYFL/SlMBP18), and FUL1 could also weakly interact with EJ2 (Supplemental Figure 1). Part of the interaction pairs we tested has been investigated before in a tomato MADS-domain interaction screen (Leseberg et al., 2008). We could reproduce all the previous results, except for interactions of FUL2 and MBP20 with MBP13, and FUL2 with MBP24. In conclusion, FUL2 can form most protein-protein interactions, which, together with its broad expression pattern, suggests that it can fulfil multiple functions in tomato, similar to FUL in Arabidopsis. FUL1 and MBP20 share a reduced set of interaction partners and MBP10 has the smallest set of interactors. The low number of interactors for MBP10 in combination with its weak overall expression pattern hints at relaxed selective pressure on this gene.

### Different variants of the *FUL1* gene exist in tomato cultivars

Upon cloning and sequencing of the *FUL1* cDNA of tomato cv. Moneyberg, which was used in our experiments, we noticed a deletion of the 5^th^ base (G) of the last exon as compared to the reference sequence (cv. Heinz). This deletion is predicted to result in a 205 amino acid protein, lacking the C-terminal 40 amino acids as compared to the reference (Figure 1C). Inspection of the genome sequence of 38 re-sequenced cultivated tomato varieties (Aflitos et al., 2014) showed that approximately half (17) contain this deletion (Supplemental Figure 2A). The latter included the much-studied cv. ‘Ailsa Craig’. The deletion was not detected in any of the re-sequenced wild accessions, suggesting that it may have first emerged after domestication. Although the deletion results in a C-terminally truncated protein (FUL1-ΔC), no differences in interactions were observed in a yeast two-hybrid assay (Supplemental figure 2B). An Electrophoretic Mobility Shift Assay (EMSA) also revealed that FUL1-ΔC can bind to a CArG-box-containing DNA fragment as a heterotetramer with TM3 (Fig. 1D). Thus, *in vitro* DNA-binding and protein-protein interaction capacities appear normal for FUL1-ΔC, indicating, together with the already described fruit ripening function for FUL1-ΔC in the cultivars Moneyberg and Ailsa Craig (Wang et al., 2019; Bemer et al., 2012), that the truncated protein is functional. To investigate whether there could be a link between the occurrence of the *FUL1*-Δ*C* allele and certain crop traits, we examined the recently published dataset of Roohanitaziani et al. (2020), in which a wide variety of cultivars and wild species has been characterized for many traits, including pedicel abscission, flowering time, inflorescence architecture, fruit development and fruit ripening. However, we did not find significant differences in any of these traits between the cultivars with a full-length *FUL1* allele and those with a truncated allele (Supplemental figure 2C and 2D), suggesting that the occurrence of the *FUL1*-Δ*C* allele does not have a major impact on the investigated features.

### FUL2 and MBP20 promote flowering in the primary and sympodial shoots

To dissect the biological roles of the *SlFUL* genes *in planta*, we generated loss-of-function single- and higher-order mutants with the CRISPR/Cas9 method. The first coding exon of each gene was targeted with three single-guide RNAs. After stable tomato transformation of the cultivar Moneyberg, we screened several independent first-generation (T0) transgenic lines for the presence of insertion/deletion (indel) alleles by PCR and sequencing. Then we generated the progeny of the primary transgenics (T1) and selected two different homozygous indel alleles for each gene, encoding truncated proteins caused by frameshifts and premature stop codons (Supplemental Figure 3). T2 phenotyping of the selected mutants revealed that the plants with homozygous knockout alleles for either *FUL2* or *MBP20* exhibited delays in primary shoot flowering, switching to reproductive growth after 13 leaves, compared to 11 in the wild type (Figure 2A). The number of days to first flowering was also significantly decreased, but was more variable between individuals of the same genotype (Supplemental Figure 4A). No significant effect in the number of leaves was observed for the *ful1* or *mbp10* single mutants (Figure 2A, Supplemental Figure 4B). The observed delay in the transition to flowering was more pronounced in higher-order mutants, with approximately three leaves extra in *ful2 mbp20, ful1 ful2 mbp20* and *ful1 ful2 mbp10 mbp20* compared to the wild type (Figure 2A). The fact that neither the *ful1* nor the *mbp10* mutations enhanced the mutant phenotype, suggests that *FUL2* and *MBP20* are the most important *FUL*-like genes for promoting the floral transition. In addition to a delay in primary shoot transition, we also observed late flowering in the sympodial shoots of the same set of mutants, increasing to an average of four leaves per sympodial shoot, while the wild type always has three (Figures 2B and 2C, Supplemental Figure 4C and 4D). To further investigate the delayed flowering phenotype, we imaged the primary shoot meristems of the wild type and quadruple mutant at different leaf stages and determined the timing of the reproductive stage transition (Supplemental Figure 4E). In the wild type, meristem transition proceeded rapidly with a visible doming after formation of nine leaves. However, in the quadruple mutant, doming was initiated later and proceeded slower, resulting in completion of the FM only after 12 leaves. In conclusion, *FUL2* and *MBP20* additively regulate the timing of flowering in both the primary shoot and the sympodial shoots. Later during development, *ful1* single mutants also showed an increase in leaf number (Figure 2C, Supplemental Figure 4D), suggesting that *FUL1* plays a minor role as well.

**Figure 2.**
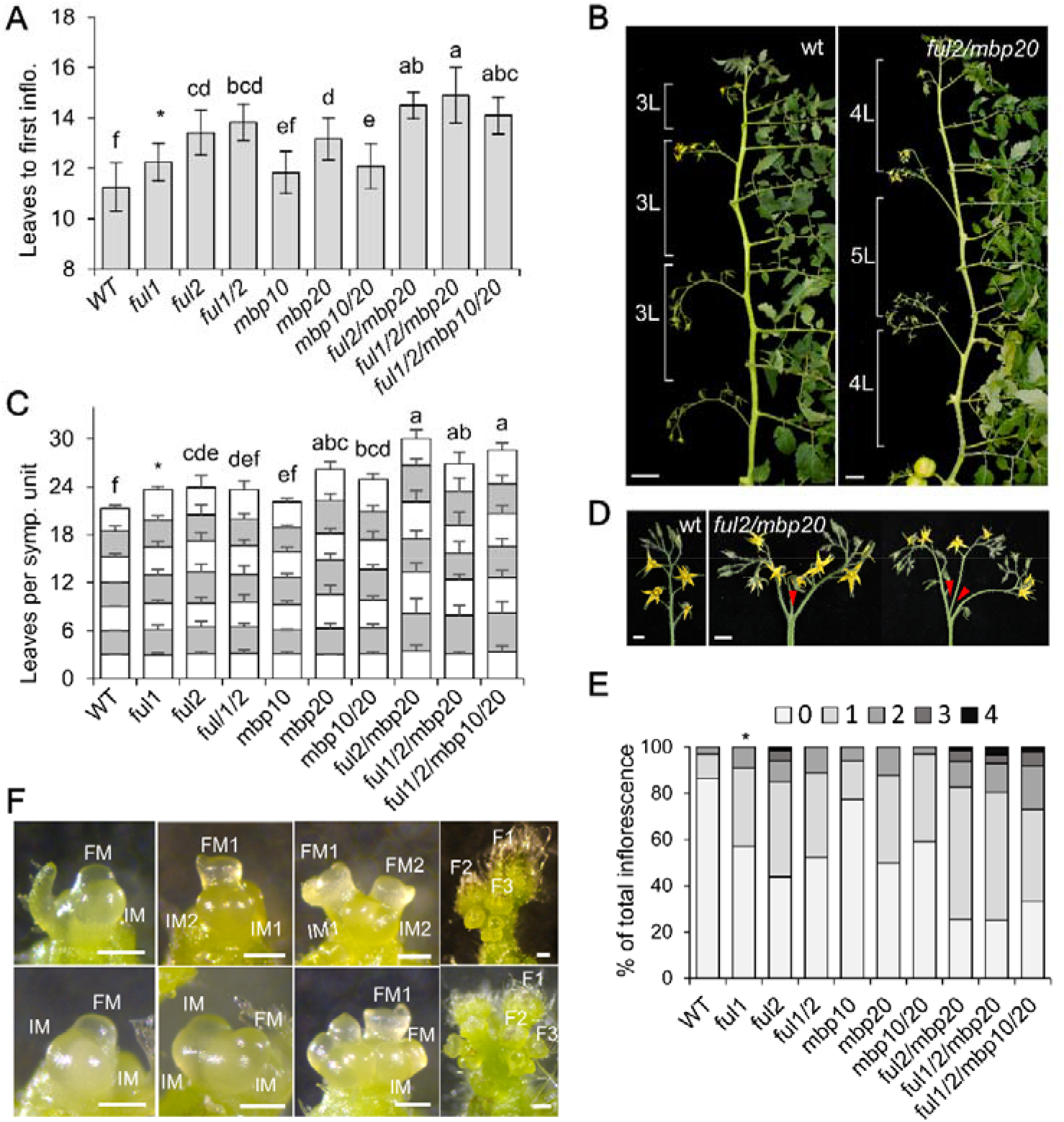
Disruption of the *FUL*-like genes results in delayed flowering and enhanced inflorescence branching. **(A)** Quantification of primary shoot flowering for wild-type (WT) and *slful* mutants. **(B)** Representative sympodial shoots from WT plants and the *ful2 mbp20* mutant. L: leaf; Scale bars: 5 cm. **(C)** Cumulative numbers of leaves in successive sympodial units for the first seven units in WT and *slful* mutants. The sum of the leaves was used to test the significance. **(D)** Representative images of wild type and mutant (branched) inflorescences. Red arrowheads indicate branching events. Scale bars: 2 cm. **(E)** Proportion of branched inflorescences in the indicated genotypes. **(F)** A developmental series of sympodial meristems of WT (upper) and quadruple mutant (lower panel) from the FM stage to an inflorescence. FM: floral meristem; IM: inflorescence meristem; F: flower. White bar: 200 μm. In A and C, mean values (± SD) were compared between genotypes using one-way ANOVA followed by a post hoc LSD test, different letters indicate the difference at P<0.05 level, six individual T2 offspring plants were analyzed per line and the data from the two different genotypes were combined for each mutant (e.g. 2×6 individuals for *ful1* etc.). The asterisk in A,C and E indicates that the *ful1* data is acquired from a second phenotyping experiment and normalized against a wild type control.

### FUL1, FUL2 and MBP20 control inflorescence architecture

We observed that the mutant plants had more branched inflorescences than wildtype plants, which typically produced only non-branched inflorescences (Figure 2D). We quantified the branching events for the first seven inflorescences of each plant in the T2 generation. With the exception of *mbp10*, all mutants showed increased inflorescence complexity, ranging from bi-parous to quintuple-parous inflorescences (Figure 2E). Notably, 13.6% of the inflorescences from wild-type plants branched, while *ful1, ful2* and *mbp20* lines produced 43.1%, 56.1% and 50% branched inflorescences, respectively. In higher-order mutants, branching increased further to ~75% in both the *ful2 mbp20* and the *ful1 ful2 mbp20* mutants. *mbp10* mutants were hardly branching, similar to the wild type, while *mbp10 mbp20* mutants were identical to *mbp20* mutants. These results indicate no additional contribution of *mbp10* or *ful1* to the branching phenotype of the mutants. Surprisingly, the *ful1* mutant did exhibit enhanced branching, but its mutation did not further enhance the phenotype of the *ful2 mbp20* mutant, suggesting that *FUL1* function depends on *FUL2* and/or *MBP20* (Supplemental Figure 4F). Interestingly, all first inflorescences did not branch, except for the first inflorescence of *ful2 mbp20* (3 plants out of 11) and *ful1 ful2 mbp20* (2 plants out of 11). Higher order branching events (i.e. quintuple parous) were only observed in mutant lines where *ful2* was included, suggesting that *FUL2* has the most prominent role in the repression of inflorescence branching. We quantified the number of flowers on the second to fourth inflorescences. The more complex inflorescences developed more flowers, increasing from on average 10 in wild-type inflorescences to approximately 20 in the higher order mutants (Supplemental Figure 4G). To investigate whether the increased branching could also be linked to delayed floral meristem maturation, as observed for the *j2 ej2* (Soyk et al., 2017), *s* and *an* mutants (Lippmann et al., 2008), we examined different stages of meristem development under the microscope (Figure 2F). We observed delayed FM development of the sympodial shoots, allowing the formation of a second IM before FM determination. The frequency of additional IMs was variable, but most similar to that of the *j2 ej2* mutant (Soyk et al., 2017). In conclusion, mutations in *FUL2* and *MBP20*, individually or combined, result in increased branching during inflorescence development. Both genes thus regulate inflorescence architecture in an additive manner, probably by regulating FM determination. *FUL1* is also involved in this process as is shown by the *ful1* single mutant phenotype, but its role is masked in higher order mutants that contain *ful2* and *mbp20* alleles.

### *MBP10/20* do not contribute to fruit development and ripening

*FUL1* and *FUL2* were reported as redundant regulators of tomato fruit ripening, and *FUL2* has an additional function in early fruit development (Bemer et al., 2012; Wang et al., 2019). Since *MBP20* is weakly expressed in carpels and early stages of fruit development (Figure 1A), we asked whether it could function in fruit development as well. We therefore examined fruit development and ripening in the different mutant lines. As reported before, the *ful2* mutant fruits were smaller with stripes on the pericarp, while the *ful1 ful2* mutant fruits were more severely impaired in ripening (Wang et al., 2019). *ful2 mbp20* mutant fruits had the same phenotype as *ful2* fruits, while triple (*ful1 ful2 mbp20*) and quadruple mutant fruits resembled *ful1 ful2* fruits in terms of width, Brix value, number of locules, pericarp stripes and overall external and internal appearance, indicating that *MBP10/20* do not contribute to fruit development and ripening (Supplemental Figures 5 and 6). Remarkable was the high Brix values of fruits that contained *ful2* mutant alleles, although this may to a large extent be due to the smaller size of *ful2* fruits (Supplemental Figure 5B). Interestingly, the number of locules was slightly, but significantly, enhanced in *ful2* single mutants, and in most mutant combinations that contained *ful2* (Supplemental Figure 5C and 5D), suggesting that *FUL2* is also involved in determining floral meristem size, and thereby carpel number.

### Dynamic expression of MADS-box genes in the inflorescence meristem

The CRISPR mutant analysis revealed that *FUL2* and *MBP20* promote the transition from vegetative to reproductive development and control inflorescence architecture. To further unveil the role of the tomato *FUL*-like genes in flowering, we conducted RNA-seq to compare the transcriptome dynamics during three consecutive stages (VM, TM, FM) of meristem development between the wild type and quadruple mutant. For each stage, over 30 meristems from a batch of plants were dissected and pooled for RNA extraction. Three independent batches were grown in the greenhouse at different time points to serve as biological replicates. For practical reasons, the FM and flanking IM were harvested together (further referred to as FM) (see Figure 3A). High-throughput sequencing yielded a minimum of 30M reads per sample. A PCA plot was generated of all 18 samples, which showed clear separation of the VM, TM and FM samples, although there was quite some distance between the individual TM samples, probably reflecting the transient nature of this stage (Supplemental Figure 6A). We first determined the expression of the *FUL*-like genes in the different stages of meristem maturation, revealing dynamic expression changes through the vegetative-to-reproductive transition for *FUL1, FUL2* and *MBP20* (Figure 3B). *FUL2* and *MBP20* are already expressed at the VM stage, but their expression highly increased in the TM stage. *FUL1* is more weakly expressed in the VM, but also reaches high expression levels in the TM and FM. The higher expression of *FUL2* and *MBP20* in the VM stage is in line with their prominent role in the determination of flowering time. Of the MADS-box genes encoding SlFUL interactors, *J* (the *SVP*-homolog), was highly expressed in all three stages, while the expression of the *SEP*-like genes *EJ2*, *TM29* and *J2* gradually increased from practically absent in VM to clearly expressed in FM. The *SOC1*-homologs *TM3* and *STM3* were also expressed in all three stages (Figure 3C), but their expression decreased in the FM in contrast to that of *EJ2*, *TM29* and *J2*. The other potential SlFUL interactors were only weakly expressed. To validate the RNA-seq data analysis, we confirmed the expression patterns of the *FUL*-like genes, and the genes encoding putative interactors, with qRT-PCR analysis on pooled meristem samples from independently grown batches (Supplemental Figure 7C and 7D).

**Figure 3.**
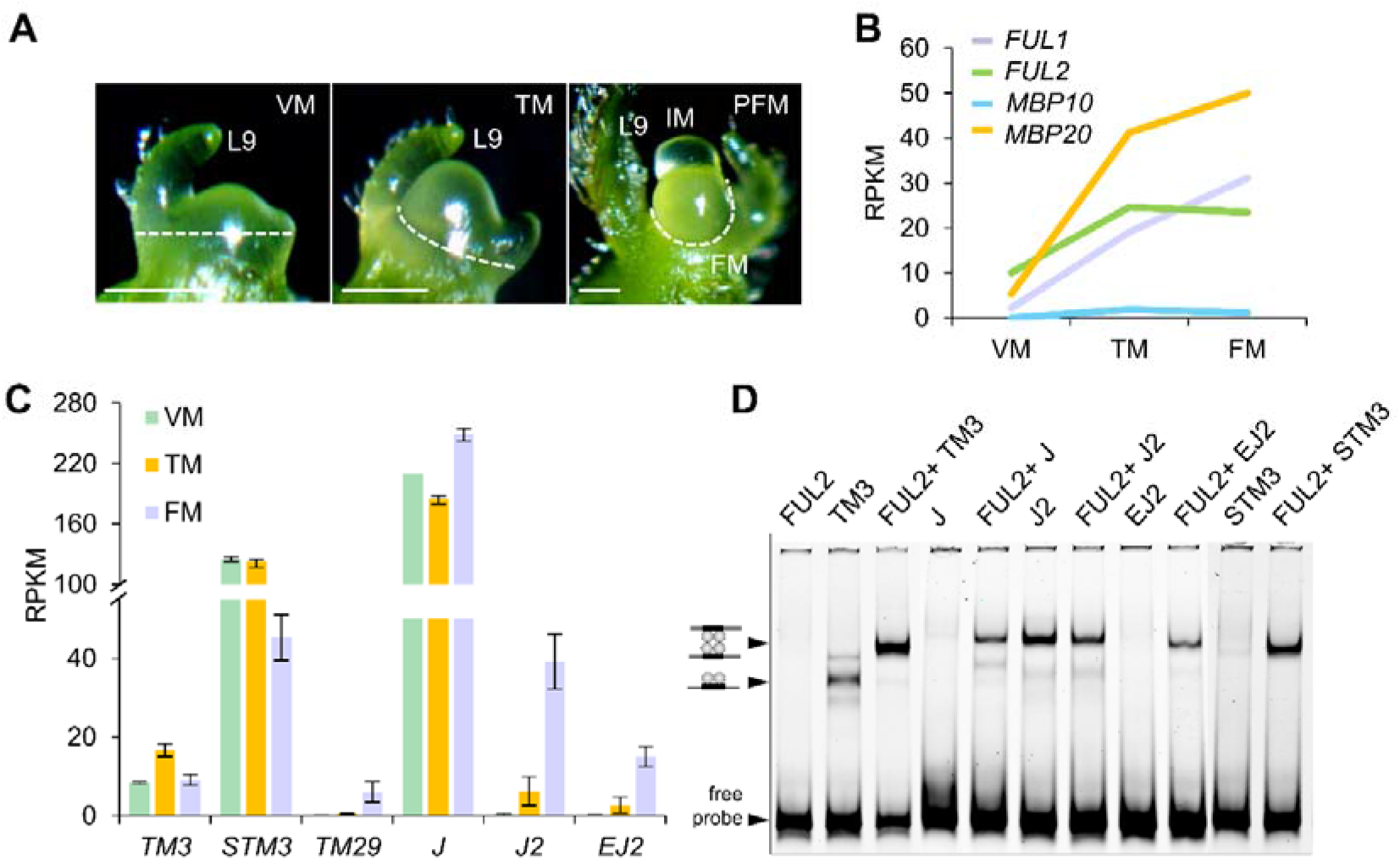
Gene expression dynamics in the primary shoot meristem. **(A)** Manual microdissection of the three successive meristem stages of primary shoot meristems for transcriptome profiling. Dashed lines indicate the dissected tissues. White bar: 100 μm. PFM: primay floral meristem. **(B)**, **(C)** Normalized gene expression (RPKM) of the *FUL*-like genes and *TM3, STM3, TM29, J, J2*, and *EJ2* in wild-type meristem stages. The values shown (mean ± SD) are the average of three replicates. **(D)** EMSA showing FUL2 interactions with MADS-domain proteins. VM: vegetative meristem; TM: transition meristem; FM: floral meristem.

The combination of the expression and interaction data provides insight into the MADS-domain complexes that may act *in planta*. Similar to the *ful1 ful2 mbp20* mutants, *j2 ej2* mutants also exhibit enhanced branching, in agreement with the increasing expression of all five genes in FMs (Soyk et al., 2017; Park et al., 2012). In contrast, mutations in *J*, *TM3* and *STM3* suppress high branching phenotypes, linked to a higher expression in the IM (Figure 3C) (Park et al., 2012; Thouet et al., 2012; Alonge et al., 2020). Thus, FUL1, FUL2 and MBP20 probably form a complex with J2 and EJ2 in the FM to promote FM maturation. However, the situation is different in the VM, where *J2* and *EJ2* are not expressed, while *TM3, STM3* and *J* are abundantly present (Figure 3C). Both the *j* and the *tm3 stm3* mutants displayed delayed flowering similar to the *ful2 mbp20* mutants (Thouet et al., 2012; Alonge et al., 2020), suggesting that FUL2 and MBP20 can physically interact with TM3/STM3 and J in the VM to induce the transition to flowering. To confirm that these complexes can be formed and bind to CArG-boxes in the DNA, we performed EMSA experiments with FUL2 or MBP20 and the putative interaction partners. Because MADS-domain proteins can only bind to the DNA probe as dimers or tetramers (De Folter et al., 2005; Immink et al., 2009), a shifted probe in the assay indicates that a dimer has been formed. Since FUL2 and MBP20 do not form homodimers (Figures 1B and 3D, Supplemental Figures 1A and 7E), we could in most cases confirm the formation of heterodimeric/tetrameric complexes by the gain of a probe shift. This was only problematic for the interactors that formed strong homodimers themselves (TM3 and J2), but for TM3, the addition of FUL2 or MBP20 resulted in a clear shift towards a tetrameric complex, confirming the yeast two-hybrid data as well (Figure 3D and Supplemental Figure 7E). Thus, based on expression patterns of the genes and interaction capacity, FUL2 and MBP20 probably interact with TM3/STM3 and J in the VM to regulate flowering time, while it is plausible that FUL1, FUL2 and MBP20 form a complex with J2 and EJ2 to promote FM meristem maturation. In addition, FUL1 and FUL2 may interact with the less abundant TM29 for this purpose.

### Identification of Differentially Expressed Genes

Comparison of WT and quadruple mutant transcriptomes revealed 131 differentially expressed genes (DEGs) for the VM stage (103 up- and 28 down-regulated in the mutant), 162 for the TM stage (137 up- and 25 down-regulated), and 185 DEGs for the FM stage (162 up- and 23 down-regulated), using FDR-corrected p-value <0.05 as a threshold for significance and a Log_2_ fold change >1.0. These genes significantly overlapped between stages, with 23 genes differentially expressed in all three stages (Supplemental Figure 7B). Many more genes were upregulated in the mutant than there were down-regulated, pointing towards a general repressive function of SlFUL-containing complexes, in agreement with data from Arabidopsis FUL studies (Ferrándiz et al., 2000; Bemer et al., 2017; Balanzà et al., 2018). A large proportion of the DEGs are involved in metabolic processes, such as terpene synthesis or the phenylpropanoid pathway, but the corresponding genes were in general weakly expressed in the meristem (Supplemental Table 1). Notably, the phenylpropanoid pathway is also controlled by FUL1/2 in tomato fruits (Bemer et al., 2012), indicating that the regulation of some identified DEGs is probably of greater importance in other tissues. The DEG lists also contained several interesting genes that are possibly involved in flowering, although previously described tomato key regulators, such as *AN*, *FA*, *SFT*, *SP* and *S* were not among the DEGs (Supplemental Table 1). We searched the list of DEGs for genes that may explain the flowering phenotypes instead, and identified a few homologs of known Arabidopsis flowering genes, such as *VRN1* and *AHL15*, which are in Arabidopsis involved in the regulation of flowering time and axillary meristem outgrowth, respectively (Levy et al., 2002; Karami et al., 2020a). Also, the MADS-domain factors *TM3* and *SlMBP13* were significantly upregulated in the quadruple mutant in all three meristem stages. Most interestingly, however, is the identification of four cytokinin signaling genes as targets of the SlFULs.

### The tomato FUL-like proteins repress negative regulators of cytokinin signaling

Compelling evidence shows that cytokinin (CK) is required for SAM activity and FM initiation, and that the interplay of transcription factor regulation and CK signaling controls the size and activity of the shoot apical meristem (Kurakawa et al., 2007; Bartrina et al., 2011; Han et al., 2014). Moreover, a recent report showed that the cytokinin reporter TCSv2 is highly expressed in tomato IM/FM meristems, with a particular high signal in doming meristems (Steiner et al., 2020). In our list of DEGs, we identified several genes involved in CK signaling, namely three *CYTOKININ OXIDASEs* (*CKXs*), *CKX1/2/6*, which degrade bioactive cytokinins, and one *type-A ARABIDOPSIS RESPONSE REGULATOR* (*ARR*), *ARR16*. We verified their differential expression with qRT-PCR in independent samples (Supplemental Figure 8A). *CKX2* and *CKX6* were upregulated in the VM and TM stages of the mutant, but not in the FM stage, while *ARR16* and *CKX1* were only upregulated in the TM stage (Figure 4A). CKXs irreversibly degrade active CKs and type-A ARRs function as negative regulators of the CK response (Brownlee et al., 1975; McGaw and Horgan, 1983; D’Agostino et al., 2000). Therefore, upregulation of *CKX1/2/6* and *ARR16* in the VM and TM stages will probably result in a reduced CK content and responsiveness. We further investigated whether FUL2 and MBP20 can directly repress *CKX* and *ARR* gene expression by binding to their promoters. We therefore scanned the up- and downstream regions of the *ARR* and *CKX* genes for CArG-box motifs, the binding sites for MADS-domain proteins (Kaufmann et al., 2009; Aerts et al., 2018). Putative CArG-boxes were present in all differentially expressed *ARR* and *CKX* genes (Supplemental figure 8B). To test whether FUL2 and MBP20 can bind to these, we performed EMSAs using fragments containing these CArG-boxes as native probes. Because MADS-domain proteins need to form a dimer to bind to the DNA, we tested TM3-FUL2 and TM3-MBP20 heterodimers, as these proteins form strong heterodimers in yeast and are probably interacting in the VM/TM. In addition, as shown above (Figure 3D), the FUL2/MBP20-TM3 tetrameric complex can be easily distinguished from the TM3 homodimeric complex in EMSA assays. We detected clear shifts for all tested regulatory fragments (Figure 4B), suggesting that the FUL2-TM3 and MBP20-TM3 heterodimers can physically bind to the tested *CKX* and *ARR* genes. To investigate whether the CArG-box is essential for the binding, we also generated mutated probes, in which the CArG-box was mildly perturbed by a single-nucleotide mutation in the center of the motifs. This probe mutation abolished or reduced the binding in all cases except for *CKX1* (Figure 4B), confirming the importance of the CArG box for the binding of the heterodimers. To determine whether FUL2 and MBP20 both play a role in repressing the cytokinin signalling genes, we harvested meristems from *ful2* and *mbp20* single mutants and performed qRT-PCRs to determine the upregulation of the *CKX/ARR* genes. Upregulation was visible in both single mutants, but was in the VM stage more distinct in the *ful2* mutant than in the *mbp20* mutant, in line with the higher expression of *FUL2* at this stage (Figure 3B). In the TM stage, both mutants showed a similar mild upregulation. The upregulation in the single mutants was considerably weaker than in the quadruple mutant, reflecting the partially redundant functions of both genes. These results suggest that both FUL2 and MBP20 directly bind to the promoter of the *CKX1/2/6* and *ARR16* genes to repress their expression and thereby upregulate cytokinin signaling in the vegetative meristem at the start of the transition to flowering.

**Figure 4.**
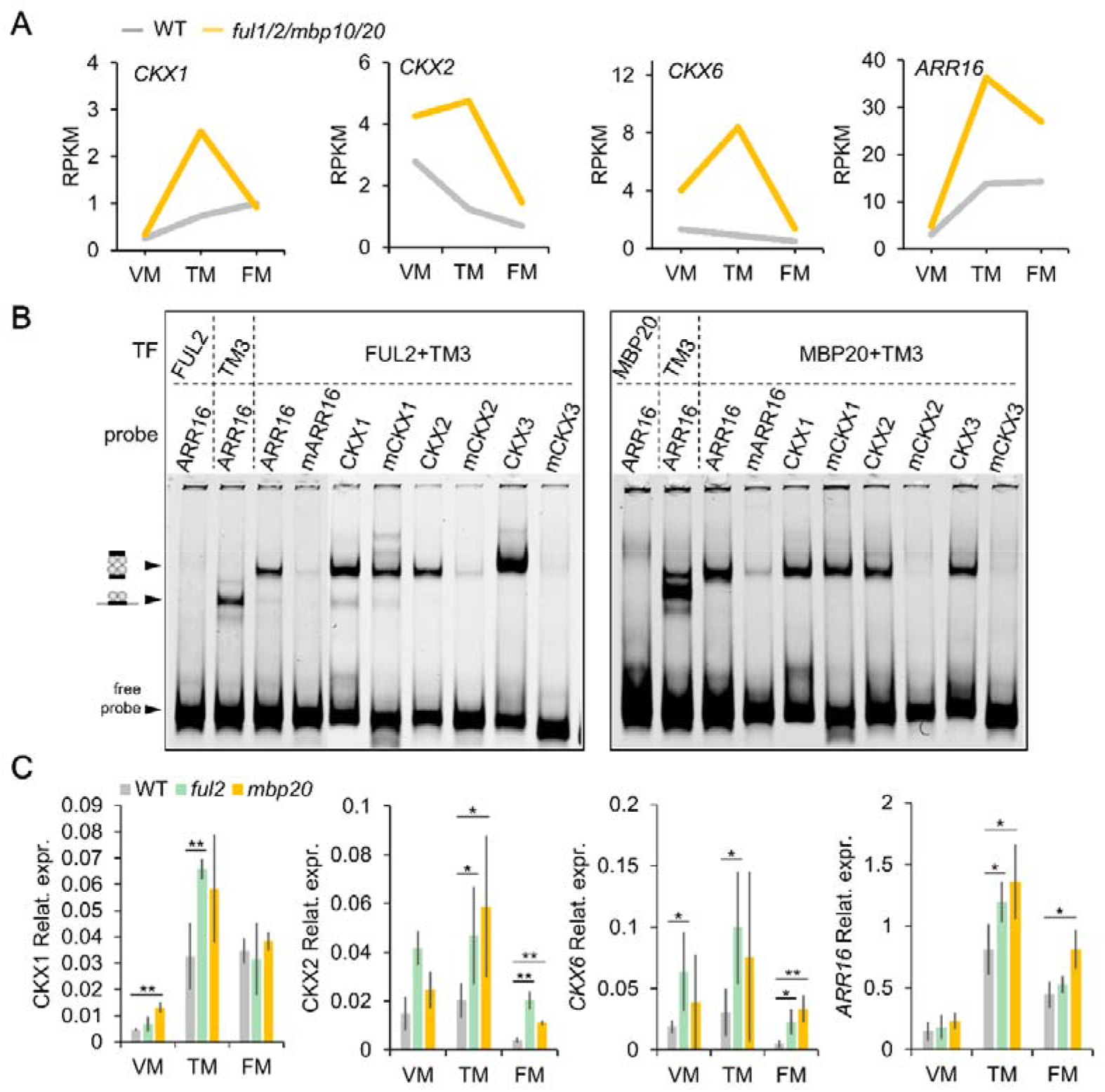
*FUL2* and *MBP20* regulate *CKX1/2/6* and *ARR16* expression. **(A)** Normalized gene expression (RPKM) for *CKX1/2/6* and *ARR16* across vegetative and reproductive meristem stages. **(B)** EMSA showing that FUL2-TM3 directly binds to promoter fragments of *CKX1/2/6* and *ARR16 in vitro*. The arrow indicates the shift of the probe caused by the binding of FUL2-TM3. The mutated versions of the promoter fragments are indicated with an ‘m’. TF: transcription factor. **(C)** Expression of *CKX1/2/6* and *ARR16* during SAM transition in wild-type, *ful2* and *mbp20* obtained by qRT-PCR. Significant differences were calculated using one-tailed Student’s t test (*, ≤ 0.05 and **, P ≤ 0.01). The values shown (mean ± SD) are the average of three replicates. VM: vegetative meristem; TM: transition meristem; FM: floral meristem.

### Transcriptome analysis of the first sympodial shoot FM reveals genes possibly involved in the branching phenotype

Searching for DEGs possibly involved in the flowering phenotype of the quadruple mutant, we found very few genes that could be associated with the inflorescence branching phenotype. However, as mentioned above, the primary shoot FM, which was harvested for the RNA-seq, only rarely gave rise to branched inflorescences. Ubiquitous branching in the quadruple mutant was only observed from the first sympodial shoot FM (SFM) onwards. Therefore, we performed an additional RNA-seq experiment to compare the transcriptomes of WT and quadruple mutant SFMs (Supplemental Figure 9A). This experiment, with the same set-up as described above, revealed 121 differentially expressed genes (DEGs), of which 96 were upregulated and 25 downregulated in the quadruple mutant. Previously reported key regulators of branching, such as *S, FA* and *AN*, were not in the list of DEGs. The expression of the *SP* gene, which suppresses the reproductive transition of the sympodial shoot meristem (Pnueli et al., 1998; Thouet et al., 2008), was remarkable, as it varied considerably between samples. (Supplemental Figure 8B). To identify genes possibly responsible for the inflorescence branching in SFMs, we searched for flowering-related genes that were differentially regulated in the SFM samples, but not in the FM samples (Figure 5A). Four transcription factors were identified that may be involved in the regulation of SFM maturation: *APETALA 2b* (*AP2b*) (Karlova et al., 2011), *AP2c* and the MADS-domain TFs *AGAMOUS-LIKE 6* (*AGL6*) and *TM29* (Figure 5B). For *TM29, AP2b* and *AP2c*, the differential expression was confirmed with qRT-PCR on independent samples (Supplemental Figure 9C). *AP2*-like genes are angiosperm-wide regulators of both meristem development and flowering, controlling for example stem-cell maintenance in the Arabidopsis SAM (Würschum et al., 2006) and floral and spikelet meristem initiation/termination in maize (Chuck et al., 2008) in addition to their ‘floral’ roles in sepal/petal development and repression of the C-function (Yant et al., 2010; Morel et al., 2017). The *SEP*-like MADS-box gene *TM29*, which is upregulated in FMs (Figure 3C), has been associated with FM identity control (Ampomah-Dwamena et al., 2002). In addition to these specifically differentially expressed genes, two other genes that are also upregulated in the primary shoot FM, but to a lesser extent (Supplemental Table 1), are probably candidates to explain the branching phenotype as well. Mutation of the first one, TM3, results in reduced inflorescence branching in the *ej2 j2* mutant background (Alonge et al., 2020), implying that higher expression of TM3 will cause enhanced branching. The other gene is a close homolog of Arabidopsis *AHL15*, which suppresses axillary meristem maturation in Arabidopsis (Karami et al., 2020b). If tomato *AHL15* is also repressing meristem maturation, this could also contribute to the enhanced branching phenotype. In conclusion, FUL1, FUL2 and MBP20 do not seem to regulate inflorescence branching by modifying the expression of the key regulators *S*, *FA* or *AN*, but we identified several other downstream transcription factors that may be involved.

**Figure 5.**
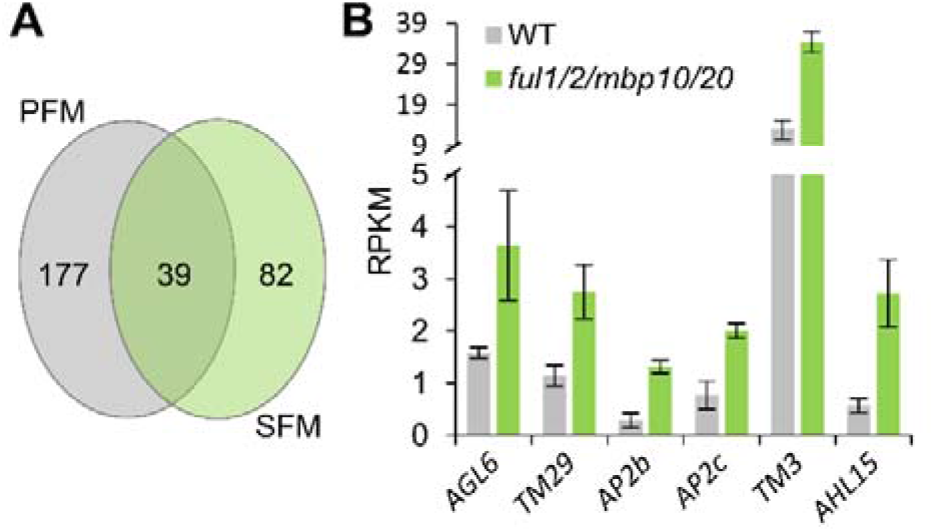
Gene expression in floral meristems of the sympodial shoot in WT and *slful* quadruple mutants. **(A)** Venn diagram showing the overlap of differentially expressed genes (DEGs) in the floral meristem of primary shoot (PFM) and sympodial shoot (SFM) of wild-type (WT) and the quadruple mutant. **(B)** Normalized gene expression (RPKM) of DEGs of interest in WT SFM. The values shown (mean ± SD) are the average of three biological replicates.

### *FUL1* expression is regulated by FUL2 and MBP20

Despite the high expression of *FUL1* in the TM and FM and the branching phenotype in the *ful1* single mutants, the *ful1* mutation does not enhance the branching phenotype of the *ful2 mbp20* mutants (Figure 2E, Supplemental Figure 4F). The distinct down-regulation of *FUL1* in the quadruple mutant may explain this apparent discrepancy (Figure 6A, Supplemental Figure 10A), and indicates that the gene is induced by FUL2, MBP20 and/or by itself via a positive (auto-)regulatory loop. Because the expression of *FUL1* is low in the VM, FUL2- and/or MBP20-containing complexes may need to bind to the CArG-boxes in the *FUL1* regulatory region to upregulate its expression in TM and FM. To test this and determine the separate effects of FUL1, FUL2 and MBP20 on *FUL1* regulation, we performed qRT-PCRs in the corresponding single mutants (Figure 6B). Downregulation was observed in all three single mutants, particularly in the FM stage, but the transcript reduction was most severe in the *ful1* mutants. The lower *FUL1* mRNA level in the *ful1* mutant may be caused by nonsense-mediated mRNA decay (NMD) as a result of the premature stop codon. However, it could also be the result of abolished FUL1 autoregulation, or a combination of both decay and disturbed autoregulation. At this point, we cannot discriminate between these possibilities. It is clear however, that both FUL2 and MBP20 positively regulate *FUL1* expression. There are four CArG-boxes in the upstream region of *FUL1* that can probably be bound by MADS-domain complexes (Supplemental Figure 10B). To test whether FUL2 and MBP20 can bind, we performed EMSAs with TM3-FUL2 and TM3-MBP20 dimers and observed clear binding to the CArG-box containing probes (Figure 6C), suggesting that *FUL1* depends on FUL2 and/or MBP20 for maximal expression in the TM and FM stages.

**Figure 6.**
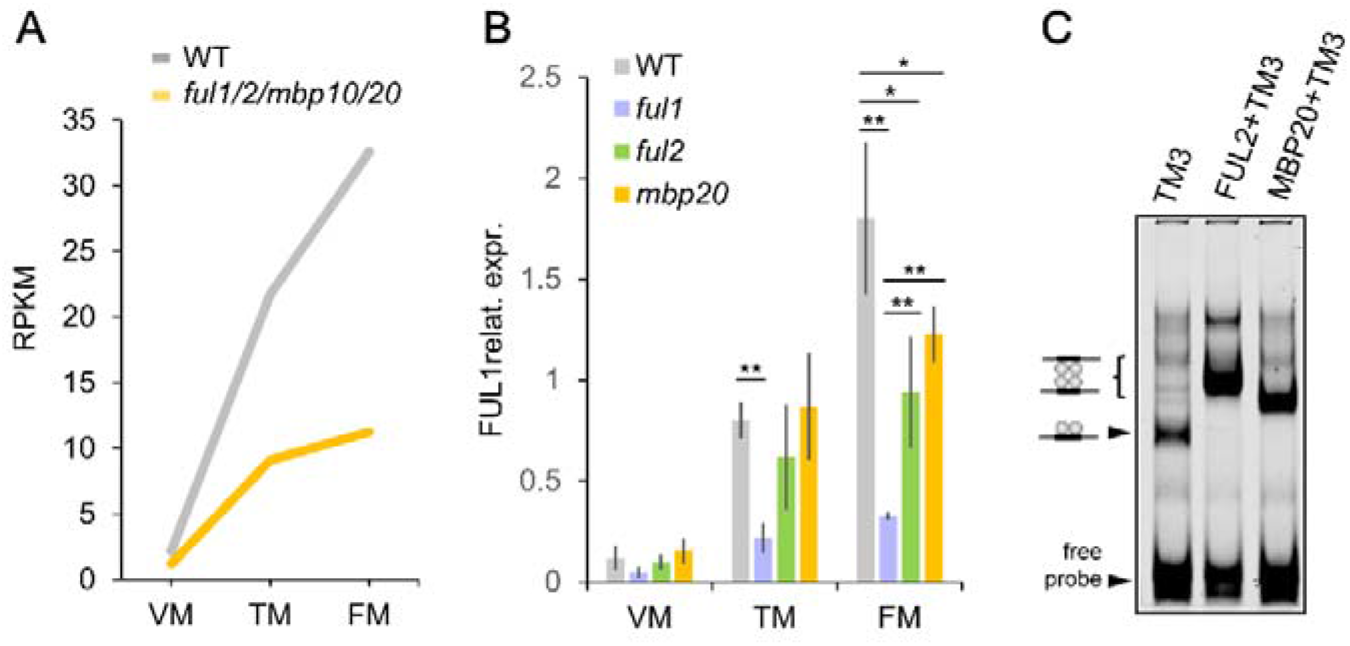
FUL2 and MBP20 regulate *FUL1* expression. **(A)** Normalized gene expression (RPKM) of *FUL1* across vegetative and reproductive meristem stages in wild type (WT) and quadruple mutant. **(B)** FUL1 expression during SAM transition in WT, *ful1, ful2* and *mbp20* tested by qRT-PCR**. (C)** EMSA showing that FUL2/TM3 and MBP20/TM3 directly bind to the promoters of *FUL1 in vitro*. Average and standard deviation of two independent replicates per stage. The arrow indicates the shift of the probe caused by the binding of FUL2-TM3 or MBP20/TM3. The values shown (mean ± SD) are the average of three biological replicates. Significant differences were calculated using one-tailed Student’s t test (*, ≤ 0.05 and **, P ≤ 0.01). VM: vegetative meristem; TM: transition meristem; FM: floral meristem.

## DISCUSSION

### Subfunctionalization of the tomato *FUL*-like genes

Following segmental or whole-genome duplication events, genes with new molecular functions can arise through sub- or neofunctionalization, resulting in divergence of biological functions. We show here that functional divergence also occurred for the tomato *FUL*-like genes after their multiplication early in the Solanaceae lineage. In addition to their previously described roles in fruit development and ripening (Bemer et al., 2012; Wang et al., 2019), we unveil that *FUL1* and *FUL2* both regulate IM development, albeit at different levels. *MBP20* promotes IM development together with *FUL2*, but does not contribute to fruit development. We did not observe any phenotype for the *mbp10* mutant, nor did the mutation enhance the phenotype in higher-order mutants. This suggests, together with its weak overall expression pattern and low number of protein-protein interactions, that *MBP10* may become a pseudogene. In line with this, *MBP10* lacks regulatory sequences in its first intron (Maheepala et al., 2019), and has a three amino-acid mutation in the I-domain, a region important for protein-protein interactions (Van Dijk et al., 2010). The loss of *MBP10* in other Solanaceae genera such as *Petunia* also hints in this direction (Maheepala et al., 2019).

Although previous overexpression studies suggested that *MBP20* functions in leaf development and *FUL2* in stem development and secondary growth (Burko et al., 2013; Wang et al., 2014a; Shalit-Kaneh et al., 2019), we did not observe aberrant phenotypes in these tissues in our knockout mutants. The most probable explanation for this discrepancy is the use of the Cauliflower 35S promoter in the previous experiments (Wang et al., 2014a; Shalit-Kaneh et al., 2019), resulting in ectopic expression and mis-regulation of target genes at a position where *FUL2* and *MBP20* are usually not expressed. Overexpressing MADS-domain proteins or dominant-negative forms of MADS proteins can also perturb complexes that involve their interaction partners, and the phenotype may thus reflect the phenotype of mutants impaired in other, interacting tomato MADS proteins. However, another possibility is that *FUL2* and/or *MBP20* function redundantly with other MADS proteins in the investigated tissues. Arabidopsis *FUL* and *SOC1* act redundantly in the regulation of secondary growth (Melzer et al., 2008), and *FUL2* may thus function redundantly with (*S*)*TM3* in the tomato stem as well. In conclusion, the four *FUL*-like genes in tomato underwent a functional divergence during evolution, but together retained functions in both inflorescence and fruit development. It is possible that some functions have remained unidentified due to redundancy with other MADS-box genes.

### The position of *FUL1* in the flower regulatory network

*FUL1* appears to act differently from *FUL2* and *MBP20* in the inflorescence meristem. It is only weakly expressed in the VM, and its high expression in TM and FM probably depends on FUL2 and MBP20, which are already expressed earlier in the VM and can bind to the *FUL1* promoter. (Auto-)regulatory loops are a common phenomenon in MADS-box gene regulation. For example, Arabidopsis *AP1* contains a CArG-box in its promoter, which can be bound by its own protein as well as by its paralog (CAL) to achieve high expression levels throughout different stages. (Ye et al., 2016). Because of the delayed induction, *FUL1* does not regulate flowering time, but does contribute to meristem maturation and thereby the repression of inflorescence branching.

Interestingly, we found that FUL1 has a premature stop codon at the C-terminus in the cultivar Moneyberg and many other cultivars. Although this truncation does not alter *in vivo* dimer formation with other MADS-domain proteins (Supplemental Figure 2B), nor disturbs tetramer formation and DNA binding (Figure 1D), the C-terminus may be important for protein activity. It contains the highly conserved, but uncharacterized, FUL-specific PQWML motif (Litt and Irish, 2003). Arabidopsis *ful* mutants complemented with a FUL copy with a mutation in this motif, were less able to rescue the silique phenotype than those transformed with a wild-type copy, suggesting that the motif is important for protein activity (McCarthy et al., 2015). Interestingly, the truncated allele has not been observed in wild relatives of tomato, and so probably first occurred after domestication (Supplemental Figure 2A). However we did not find a correlation between the presence of the *FUL1*-Δ*C* allele and any trait characterized by Roohanitaziani et al. (2020), but we cannot exclude that the allele has been selected during breeding, for example by conferring slightly larger inflorescences without severe branching.

### The role of *FUL2* and *MBP20* in the tomato flowering network

Several previously identified tomato flowering genes were revealed to be functional homologs of Arabidopsis flowering genes, such as *SFT* (*FT*) and *FA* (*LFY*) (Molinero□Rosales et al., 1999; Molinero-Rosales et al., 2004; Lifschitz et al., 2006), indicating that at least part of the Arabidopsis flowering network is conserved in tomato. However, the knowledge of the regulatory network underlying the tomato sympodial flowering pathway is still fragmented and it is yet unclear whether homologs of many important players in Arabidopsis, such as *SOC1* and *FLC*, are important for tomato flowering as well. Here, we show that tomato *FUL*-like genes regulate flowering and inflorescence development in tomato, thereby adding a piece to the tomato flowering network puzzle. In Arabidopsis, *FUL* is a target of FLOWERING LOCUS D (FD) and SQUAMOSA PROMOTER BINDING PROTEIN-LIKE (SPL) proteins in the photoperiod pathway and the age pathway (Kardailsky et al., 1999; Wang et al., 2009; Jung et al., 2016), and functions partially redundantly with *AP1* in the promotion of flowering (Ferrándiz et al., 2000). We demonstrate here that *FUL2* and *MBP20* additively promote flowering similar to their homolog in Arabidopsis, but it is yet unclear whether they act downstream of SFT and the tomato SPLs as well. However, we did identify putative SPL and FT/FD binding sites *in silico* in both the *FUL2* and *MBP20* promoter sequences, suggesting that their expression may be regulated in a similar way. Within our set of DEGs, we did not identify any of the previously identified flowering regulators (e.g. FA, S, SFT, SP), further indicating that the tomato *FUL* homologs may act downstream of, or parallel to, these factors.

MADS-domain transcription factors bind to the DNA as dimers (De Folter et al., 2005), and since the tomato FUL-like proteins cannot form homodimers, they need to heterodimerize with other MADS-domain proteins to regulate target gene expression. For the regulation of flowering time, FUL2 and MBP20 probably form a complex with TM3, STM3 and J, because the corresponding genes are highly expressed in the vegetative and transition meristem, and both *tm3 stm3* and *j* mutants display a small delay in flowering time (Szymkowiak and Irish, 2006; Alonge et al., 2020) similar to *ful2* and *mbp20*.

Downstream of the *FUL*-like genes, we discovered several repressors of the cytokinin signaling pathway that are upregulated in the VM and TM stages of the quadruple mutant, probably resulting in reduced CK levels and signaling. In tomato, the switch from vegetative to transition meristem is accompanied by cell division in the meristem, which results in the doming of the TM. Steiner et al. (2020) showed that this doming is accompanied by a high cytokinin signal in the meristem. Because CK is a prominent inducer of cell proliferation (Miller et al., 1955), it can probably positively regulate cell division during SAM maturation to allow doming of the meristem and the transition to flowering. In line with this hypothesis, the reduced CK levels may inhibit SAM doming and thereby delay flowering. In Arabidopsis, CK deficiency through overexpression of *CKXs* diminishes shoot meristem activity and indeed retards flowering (Werner et al., 2003). In addition, initiation of both the axillary meristem and the FM revealed to require a cytokinin signaling pulse (Han et al., 2014; Wang et al., 2014b). Our data suggest that FUL2 and MBP20 may promote flowering through indirect regulation of CK levels and responsiveness by directly repressing *CKX* and *type-A ARR* expression, respectively (Figure 7).

**Figure 7.**
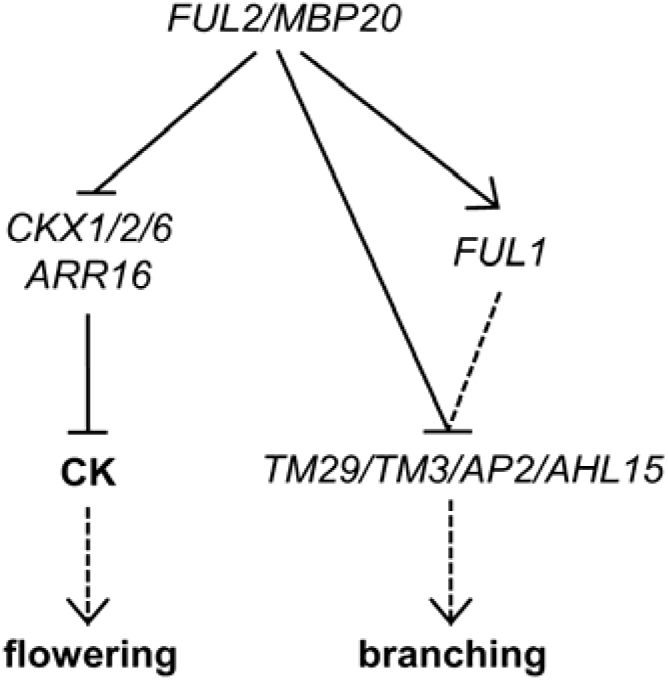
Model of SIFUL regulation of flowering time and inflorescence branching in tomato. The connections between the different regulators are based on the results of this study and other work described in the text. Solid lines display confirmed interactions, while dashed lines represent putative interactions. CK is cytokinin.

In other species, such as Arabidopsis and Petunia, *AP1/FUL*-like genes are involved in the establishment of IM/FM identity, and combinatorial mutations result in a loss of identity, leading to a non-flowering phenotype. The Arabidopsis *ap1 cal* double mutant forms only IMs, because FM identity is lost, and mutation of *ful* also impairs IM identity, resulting in a more severe vegetative phenotype (Ferrándiz et al., 2000). In Petunia, the four FUL/AP1 genes appear to function redundantly in the establishment of IM/FM identity, and higher order mutants remain for a long time in the vegetative stage (Morel et al., 2019). In tomato, vegetative reversion occurs after a few flowers have formed in mutants of the *AP1*-homolog *MC* and the *SVP*-homolog *J*, suggesting that these are required for FM fate as well (Szymkowiak and Irish, 2006; Thouet et al., 2012). We observed this phenotype only occasionally in the *ful* mutants, suggesting that the promotion of IM fate is mainly regulated by *J* and *MC*. It is possible however, that the tomato *FUL*-like genes act redundantly with *MC*, which would reflect the situation in Petunia.

### The role of the *SIFULs* in the regulation of inflorescence architecture

We show that loss of function of *FUL1, FUL2* and/or *MBP20* results in a branched inflorescence that produces an increased number of flowers. The phenotype is variable however, with some mutant inflorescences staying single-parous, while others form up to five branches. Our transcriptome analysis revealed that this branching is not caused by regulation of *S*, *AN* or *FA*, indicating that they function upstream of, or in parallel with, the SlFUL-containing complexes. Remarkable was the varying expression of the TFL-ortholog *SP*, which may be involved in branching by the maintenance of IM identity. *SP* seemed higher expressed in the quadruple mutant, but this effect was not significant due to the large variation (Supplemental Figure 7B). However, because the branching phenotype is also highly variable, we cannot exclude that *SP* is somewhat regulated by the tomato FULs in IM/FM, thereby exerting an effect on the branching phenotype. This would be similar to the repression of the legume *TFL*-homolog by the euFULII clade proteins VEG1 (pea) or MtFUL1-c (Medicago) (Berbel et al., 2012; Benlloch et al., 2015; Cheng et al., 2018; Zhang et al., 2021).

The branching phenotype of the quadruple mutant is similar to that of the *j2 ej2* mutant (Soyk et al., 2017) and our microscopic analysis suggests that it is caused by delayed maturation of the FM as well. Given our *in vitro* interaction data, which show that the FULs can interact with J2 and EJ2, it is plausible that they act together in a complex to promote FM maturation and suppress inflorescence branching. Both the *FULs* and *J2/EJ2* are clearly expressed in the FM in our data (Figure 3C), although our sampling method did not allow a clear distinction between FM and flanking SIM. However, the data from Park et al., (2012) are based on FMs that were completely isolated, and they describe high expression for *FUL1* and *FUL2* in the FM, while Soyk et al. (2017) describe the same for *J2* and *EJ2*, in agreement with an important role of a FUL1/FUL2/MBP20-J2/EJ2 complex in the regulation of FM maturation. Genetic experiments revealed that the *j2 ej2* mutant phenotype is additive to that of *s*, indicating that *J2* and *EJ2* function separately from the *S* gene (Soyk et al., 2017), and the same probably accounts for the SlFULs.

Our transcriptome analysis of the SFMs unveiled several genes encoding transcription factors that could function in the regulation of FM maturation downstream of the SlFULs (see Figure 7). Two of these, *TM29* and *TM3*, are also encoding for MADS-domain transcription factors, which are upregulated in the quadruple mutant FMs. In particular TM3 is interesting, because its expression is high in VM and TM, but drops in FM. This suggests that TM3 is repressing FM maturation, in line with the observation that the *tm3 stm3* double mutants represses the enhanced branching phenotype of *j2 ej2*. The upregulation of TM3 will thus delay FM maturation and thereby enhance branching. Indications for the involvement of the other genes, *AHL15, AP2b and AP2c*, rather comes from research in Arabidopsis and maize. In Arabidopsis, *AHL15* is repressing meristem maturation in the axillary buds (Karami et al., 2020a), while *AP2*-like genes regulate meristem development in both Arabidopsis and maize (Würschum et al., 2006; Chuck et al., 2008). The role of *AP2b* and *AP2c* could also be linked to their role as A-class floral organ genes, because *AP2*-like genes are in many species involved in the repression of the C-function (Morel et al., 2017), of which the onset marks complete FM maturation. Which of these downstream factors is most important for the increased branching phenotype still needs to be determined with future genetic experiments and localization studies to determine what their function is in either the FM or the IM.

## MATERIALS AND METHODS

### Accession numbers

FUL1, Solyc06g069430; FUL2, Solyc03g114830; MBP10, Solyc02g065730; MBP20, Solyc02g089210; J, Solyc11g010570; J2, Solyc12g038510; EJ2, Solyc03g114840; TM29, Solyc02g089200; MADS-RIN, Solyc05g012020; TM5, Solyc05g015750; TM3, Solyc01g093965; STM3, Solyc01g092950; SlMBP18, Solyc03g006830; TAG1, Solyc02g071730; SlMBP24, Solyc01g105800; SlMBP13, Solyc08g080100; SlMBP14, Solyc12g056460; SlMBP9, Solyc04g076680; SlMBP12, Solyc12g088090; SlMBP22, Solyc11g005120; MADS-MC, Solyc05g056620; AHL15, Solyc12g087950; AP2b, Solyc02g064960; AP2c, Solyc02g093150; AGL6, Solyc01g093960; CKX1, Solyc04g016430; CKX2, Solyc10g017990; CKX6, Solyc12g008900; ARR16, Solyc06g048930; AP2b, Solyc02g064960; Ap2c, Solyc02g093150. Supplementary Table 1 contains the accession numbers of the DEGs.

### Plant materials and growing conditions

Tomato cv. Moneyberg was used for the *Agrobacterium tumefaciens-mediated* transformation experiments (Van Roekel et al., 1993). Tissue culture was conducted in a growth chamber with 16 h light and 8 h dark at 25 °C. After rooting, the transformed plants were cultivated in a 21 °C growth chamber (16h light/8h dark). 25-day old plants were moved to the greenhouse and grown under ambient temperatures and natural light, supplemented with artificial sodium lights.

### qRT-PCR analysis

For qRT-PCR analysis of *SlFUL* expression, root, shoot, leaves, flower organs, and fruits of different stages were harvested from wild-type tomato plants. RNA was extracted with a CTAB/LiCl method (Porebski et al., 1997), DNase treated with Ambion Turbo DNase (AM1907) and cDNA was synthesized with the iScript cDNA synthesis kit (Bio-Rad). Real-time RT-PCR was performed with the iQ SYBR Green Supermix from Bio-Rad with a standard 2-step program of 40 cycles, annealing at 60 °C. Primer efficiencies were tested beforehand and only primer pairs with equal efficiencies were compared (all primer sequences, including reference primers, are listed in Supplemental Table 2).

### Yeast two-hybrid

Protein-protein interaction assays in yeast were performed using the GAL4 System using Gateway vectors as described (De Folter and Immink, 2011). The coding sequences for bait proteins and prey proteins were cloned into the pDEST32 and pDEST22 vectors respectively, and the vectors were transformed into the PJ69-4A and PJ69-4α yeast strains. The interaction screen was performed using-LWH dropout medium, supplemented with 3 mM 3-amino-1,2,4-triazole (3-AT) or -LWA dropout medium. Plates were incubated for 5 days at RT. All primer sequences used for cloning are listed in Supplemental Table 2.

### CRISPR construct generation and stable tomato transformation

The *ful2* and *ful1/2* transgenic CRISPR lines have been previously generated (Wang et al., 2019). The constructs for all other lines were generated using GoldenGate cloning and the MoClo toolkit according to (Weber et al., 2011). Briefly, each gRNA was fused to the synthetic U6 promoter as U6p::gRNA, and ligated in a Level 1 vector. Level 1 constructs pICH47732-NOSpro::NPTII::OCST, pICH47742-35S::Cas9::NOST, pICH47751-35S::GFP::ter35S, pICH47761-gRNA1, pICH47772-gRNA2, pICH47781-gRNA3 and the linker pICH41822 were cut/ligated into the level 2 vector pICSL4723 as described. After confirming the constructs, the plasmids were transformed into Agrobacterium strain C58C1. All primers are listed in Supplemental Table 2. The above constructs were introduced into tomato cv Moneyberg by *Agrobacterium tumefaciens-mediated* transformation. Homozygous T1 or T2 transgenic plants were used for phenotypic and molecular characterization.

### Meristem imaging

Shoot apices were dissected from young plants using a forceps and older leaf primordia were removed to expose meristems under the stereomicroscope. Immediately after dissection, live meristems were imaged using a euromex scientific camera.

### Meristem transcriptome profiling

The domesticated tomato (S. lycopersicum) cultivar Moneyberg and the homozygous *ful1 ful2 mbp10 mbp20* mutant generated in the Moneyberg background were used for transcriptome profiling. For each biological replicate sample, a batch of plants was grown and from each plant, the primary shoot meristem was harvested, either in the VM, TM or FM stage. For the SFM samples, the first FM from the sympodial shoot was harvested. About 60 plants were grown per batch (to harvest >30 meristems). All stages were harvested in triplicate for both the wild type and quadruple mutant plants. The batches for the different replicates were grown in the greenhouse sequentially. Meristems were dissected using a stereoscope, and tissue was processed for RNA stabilization using an acetone fixation technique (Park et al., 2012). RNA was extracted using the PicoPure RNA Extraction kit (Arcturus). More than 30 meristems were collected for each sample, yielding 1~3 μg RNA, which was enriched for mRNA and processed into cDNA libraries using the Illumina TruSeq Stranded Total RNA LT Sample Prep Kit (Illumina). After quality control (Qubit and Fragment Analyzer), samples were sequenced using Illumina NovaSeq 2×150 nt Paired End sequencing. Samples were randomized across sequencing flow cells and lanes within flow cells. After quality control, all data were analyzed using the CLC work package. The raw data has been deposited in GEO under accession number GSE154419. For data validation, new batches of plants were grown and processed as described above, and the samples were analysed using qRT-PCR analysis (see Supplemental Table 2 for the primers).

### Electrophoretic mobility shift assays (EMSAs)

*FUL2 and MBP20* coding sequences were amplified from wild-type Moneyberg cDNA and cloned into pSPUTK (see Supplemental Table 2 for all primer sequences). The pSPUTK promoter allowed *in vitro* protein synthesis using the TnT^®^ SP6 High-Yield Wheat Germ Protein Expression System (Promega) according to the manufacturer’s instructions. The probe fragments consisted of a region of 80-100 bp with the canonical CArG-box in the middle, and were amplified from genomic DNA; The mutated probe fragments were generated by overlapping PCR using primers that replaced one base pair in the middle of the CArG-box. EMSAs were performed essentially as described by Smaczniak *et al*. (2012) with minor modifications. Oligonucleotides were fluorescently labelled using DY-682. Labelling was performed by PCR using vector-specific DY-682-labelled primers followed by PCR purification with NucleoSpin^®^ Gel and PCR Clean-up kit (MACHEREY-NAGEL). Gel shifts were visualized using a LiCor Odyssey imaging system at 700 nm.

## ACKNOWLEDGMENTS

We would like to thank Stuart Jansma, Albert van der Veen, Janne Hageman, Siye Chen and Iris Zahn for their helpful work on yeast two-hybrid analysis, tomato transformation or meristem harvesting. We greatly appreciate Geurt Versteeg and Teus van den Brink for their help in taking care of tomato plants and collecting seeds in the greenhouse, and André Maassen and Michiel Lammers for their efforts to create enough greenhouse space. This work has been supported by a grant from the Dutch Scientific Organization (NWO) (ALWOP.199) to M.B., a fellowship from the China Scholarship Council (CSC) to X.J., a fellowship from CAPES and CAPES/Nuffic (BEX 7686/13-7) to G.L. and the Fundação de Amparo à Pesquisa do Estado de São Paulo (FAPESP process 2010/52012-4) and Coordenação de Aperfeiçoamento de Pessoal de Nível Superior (CAPES/NUFFIC-BEX 0256/13-7) to J.H.L.

## AUTHOR CONTRIBUTIONS

M.B. conceived the project and designed the experiments; X.J. performed the qPCRs, tomato CRISPR/cas9 mutagenesis, RNA-seq experiments and EMSAs; K.R, G.L. and J.H.L. did the Yeast Two Hybrid analyses; V.V. measured the fruit phenotypes; R.A.d.M. analyzed *FUL1* alleles in cultivars and assisted with the RNA-seq analysis; G.C.A. and M.B. supervised the project; M.B. and X.J analyzed the data, prepared the figures and wrote the article. All authors read and approved the final version.

## Notes

### Competing Interest Statement

The authors have declared no competing interest.

