## Supplemental Figures for "FRUITFULL-like genes regulate flowering time and inflorescence architecture in tomato"

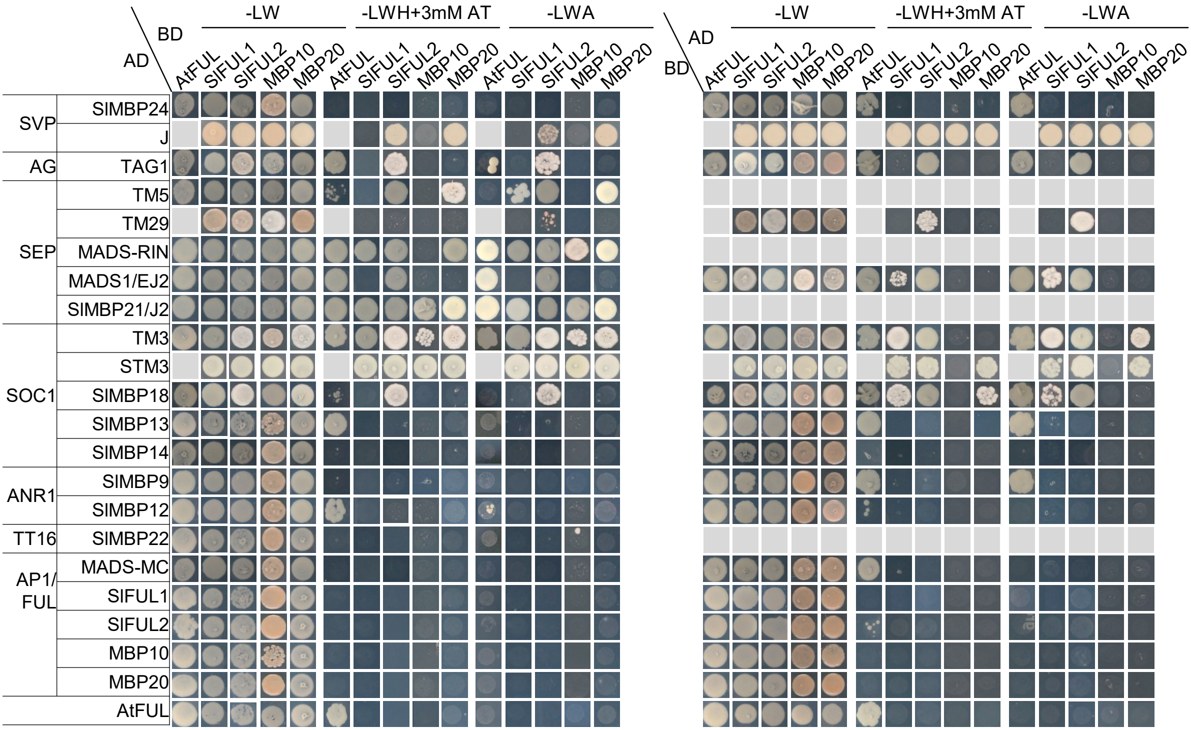

**Supplemental Figure 1. Yeast two-hybrid analysis of FRUITFULL-like proteins with MADS-box proteins from different subfamilies**. SVP: SHORT VEGETATIVE PHASE; AG: AGMOUS; SEP: SEPALATTA; SOC1: SUPPRESSOR OF OVEREXPRESSION OF CONSTANS 1; ANR1: ARABIDOPSIS NITRATE REGULATED 1; TT16: TRANSPARENT TESTA16; AP1/FUL: APETALA1/FRUITFULL. L, leucine; W, tryptophan; H, histidine; A, adenine; 3-AT, 3-amino-1,2,4-triazole. Grey boxes indicate that the interaction was not tested, in most cases because the bait gave auto-activation.

A

| Variety or Cultivar Name | Deletion present |
|---|---|
| Heinz 1706 (reference genome) | . |
| Moneymaker | X |
| Moneyberg | X |
| Ailsa Craig | X |
| Rutgers | X |
| Galina (Galina's yellow; Galina's cherry) | . |
| Ponderosa | . |
| John's Big Orange | . |
| All Round | X |
| Sonato | X |
| Cross Country | . |
| Lidi | . |
| Momatero | . |
| Large Pink | . |
| Jersey Devil | . |
| Polish Joe | . |
| Cal J TM VF | . |
| The Dutchman | . |
| Black Cherry | . |
| ANTO | X |
| Winter Tipe (nor) | X |
| Chang Li | . |
| Belmonte | X |
| Tiffen Mennonite | . |
| Wheatley's Frost Resistant | X |
| Chih-Mu-Tao-Se | X |
| ES 58 Heinz | . |
| Dolmalik | X |
| Large Red Cherry | X |
| Porter | X |
| Bloody Butcher | . |
| Brandywine | . |
| Dixy Golden Giant | . |
| Giant Belgium | X |
| Kentucky Beefsteak | . |
| Marmande VFA | X |
| Thessaloniki | X |
| Watermelon Beefsteak | . |
| S. pimpinellifolium | . |
| S. chmielewski | . |
| S. cheesmaniae | . |

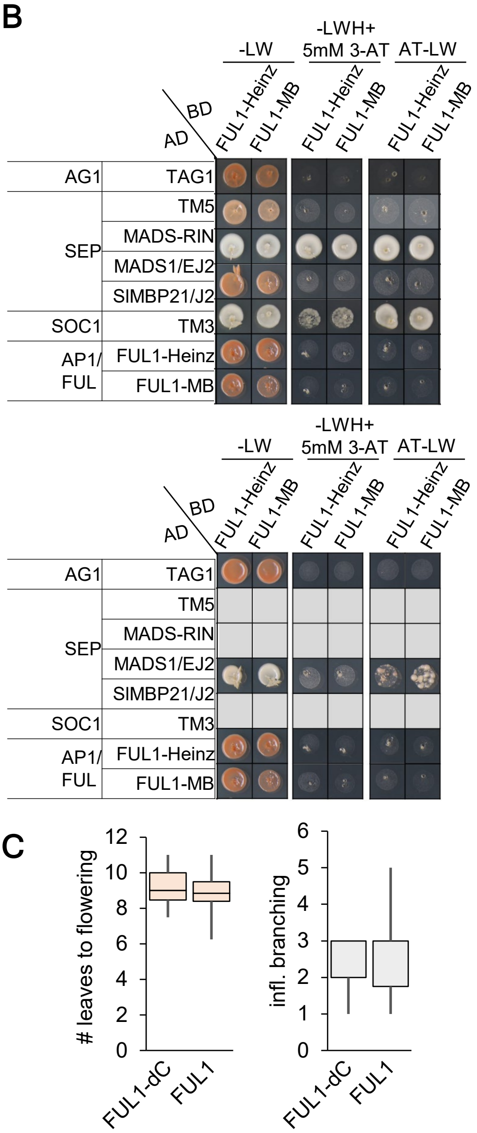

**Supplemental Figure 2. The truncated FUL1 version presents in many cultivars, but displays the same protein-protein interactions as the full-length reference protein. (A)** Table showing the occurrence of the 1nt deletion allele in 37 resequenced cultivars and wild species. X = 1nt deletion is present. **(B)** Yeast two-hybrid screening using a subset of interaction pairs and both the full-length (Heinz) and truncated (Moneyberg) FUL1 protein. **(C)** Flowering and branching phenotype of the cultivars with truncated FUL1 (FUL1-dC) and FUL1. **(D)** (see next page) Table of features grouped for cultivars with FUL1-dC and cultivars with 'normal' FUL1, according to Roohanitaziani et al. (2020).

| Genotype name | FUL allele | AZ category | Inflorensce nce branching | VOI | Flowering Time | Number of fruits | Fruit Weight (g) | Degrees Brix | Firmness (N) | Color | Shape |
|---|---|---|---|---|---|---|---|---|---|---|---|
| Moneymaker | dC | 1.0 | 2.0 | 3.0 | 8.8 | 64.0 | 41.7 | 4.9 | 52.9 | red | round |
| Ailsa Craig | dC | 1.0 | 2.0 | 1.0 | 8.8 | 66.5 | 49.6 | 4.1 | 55.4 | red | round |
| Rutgers | dC | 1.0 | 1.0 | 1.0 | 8.3 | 12.3 | 154.3 | 4.7 | 47.7 | red | Ox-heart |
| All Round | dC | 1.0 | 3.0 | 3.0 | 9.0 | 77.0 | 52.9 | 4.7 | 51.5 | red | round |
| Sonato | dC | 1.0 | 3.0 | 1.0 | 10.0 | 65.0 | 58.9 | 4.6 | 51.6 | red | round |
| ANTO | dC | 1.8 | 3.0 | 3.0 | 8.0 | 29.8 | 193.2 | 4.9 | 57.4 | red | flat |
| Winter Tipe (nor) | dC | - | 2.0 | 3.0 | 10.7 | 29.3 | 34.0 | 4.3 | 65.3 | Green | round |
| Belmonte | dC | 1.8 | 3.0 | 3.0 | 8.4 | 18.5 | 317.9 | 3.9 | 49.6 | dark pink | flat |
| Wheatley's Frost R | dC | 1.0 | - | - | 11.0 | 79.0 | 13.0 | 5.0 | 41.8 | pink | rectangular |
| Chih-Mu-Tao-Se | dC | 1.0 | 2.0 | 1.0 | 10.7 | 36.3 | 99.5 | 4.7 | 46.3 | pink | flat |
| Dolmalik | dC | 1.0 | 3.0 | 4.0 | 9.3 | 65.3 | 56.4 | 4.9 | 43.1 | red | flat |
| Large Red Cherry | dC | 1.0 | 2.0 | 4.0 | 9.0 | 106.5 | 25.3 | 5.2 | 49.3 | red | round |
| Porter | dC | 1.0 | 2.0 | 1.0 | 10.0 | 59.0 | 21.5 | 4.2 | 48.3 | pink | round |
| Giant Belgium | dC | 2.0 | 3.0 | 2.0 | 7.5 | 20.5 | 249.0 | 4.1 | 49.1 | dark pink | flat |
| Marmande VFA | dC | 1.3 | 3.0 | 1.0 | 8.5 | 35.3 | 89.8 | 4.4 | 51.0 | red | flat |
| Thessaloniki | dC | 1.0 | 3.0 | 1.0 | 9.3 | 18.8 | 127.3 | 4.7 | 39.7 | red | flat |
| | | 1.2 | 2.5 | 2.1 | 9.2 | 48.9 | 99.0 | 4.6 | 50.0 | | |
| Galina | normal | 1.0 | 1.0 | 1.0 | 9.2 | 150.3 | 13.3 | 6.0 | 53.5 | orange | round |
| Ponderosa | normal | 2.0 | 3.0 | 1.0 | 8.4 | 16.0 | 162.1 | 4.7 | 54.5 | red/orange | round |
| Ponderosa | normal | - | 3.0 | 2.0 | 8.9 | | | | 59.6 | | |
| John's big orange | normal | 1.0 | 3.0 | 1.0 | 6.3 | 32.0 | 56.6 | 4.0 | 52.8 | orange | flat round |
| Cross Country | normal | 1.0 | 1.0 | 1.0 | 9.2 | 75.5 | 36.6 | 4.2 | 37.4 | red | round |
| Lidi | normal | 1.5 | 5.0 | 5.0 | 10.1 | 459.0 | 6.2 | 6.3 | 55.4 | yellow | ovate |
| Momatero | normal | 1.0 | 2.0 | 4.0 | 10.0 | 29.3 | 150.7 | 4.8 | 55.1 | pink | flat |
| Large Pink | normal | 1.0 | 3.0 | 1.0 | 7.7 | 19.8 | 244.5 | 3.7 | 49.0 | pink | flat |
| Jersey Devil | normal | 1.0 | 3.0 | 4.0 | 11.0 | 13.8 | 105.8 | 4.7 | 53.4 | red | oxheart |
| Polish Joe | normal | 2.0 | 3.0 | 3.0 | 8.2 | 27.8 | 265.9 | 3.9 | 59.8 | pink | heart |
| Cal J TM VF | normal | 2.5 | 1.0 | 1.0 | 8.0 | 52.5 | 46.3 | 4.7 | 47.9 | red | round |
| The Dutchman | normal | 1.5 | 3.0 | 3.0 | 9.8 | 18.5 | 306.7 | 4.6 | 51.9 | pink | flat |
| Black Cherry | normal | 1.0 | 3.0 | 1.0 | 8.9 | 187.5 | 14.7 | 6.4 | 48.5 | dark pink | round |
| Chang Li "*L. esculen* | normal | 1.0 | - | - | 8.7 | 66.0 | 15.3 | 5.5 | 54.0 | yellow | round |
| Tiffen mennonite | normal | 1.0 | 3.0 | 3.0 | 8.5 | 23.3 | 142.9 | 4.7 | 56.7 | pink | flat |
| ES 58 Heinz' *L. escu* | normal | 1.0 | 1.0 | 1.0 | 9.5 | 18.5 | 99.4 | 3.5 | 41.8 | red | round |
| Bloody Butcher | normal | 1.3 | 3.0 | 4.0 | 9.5 | 50.8 | 43.8 | 5.4 | 52.8 | red | flat |
| Brandywine | normal | 1.0 | 1.0 | 1.0 | 9.5 | 14.3 | 231.4 | 4.7 | 43.3 | dark pink | round |
| Dixy Golden Giant | normal | 1.0 | 3.0 | 1.0 | 9.1 | 14.8 | 253.1 | 4.9 | 49.5 | orange | flat |
| Kentucky Beefstea | normal | 1.3 | 4.0 | 3.0 | 8.0 | 14.3 | 243.4 | 3.7 | 45.5 | orange | flat |
| Watermelon Beefs | normal | 2.0 | 3.0 | 4.0 | 8.4 | 5.8 | 305.7 | 5.1 | 51.0 | pink | ovate |
| | | 1.3 | 2.6 | 2.3 | 8.9 | 64.5 | 137.2 | 4.8 | 51.1 | | |
| **T-test** | | **0.571999** | **0.732143** | **0.956789** | **0.420505** | **0.49979** | **0.375488** | **0.430103** | **0.578271** | | |

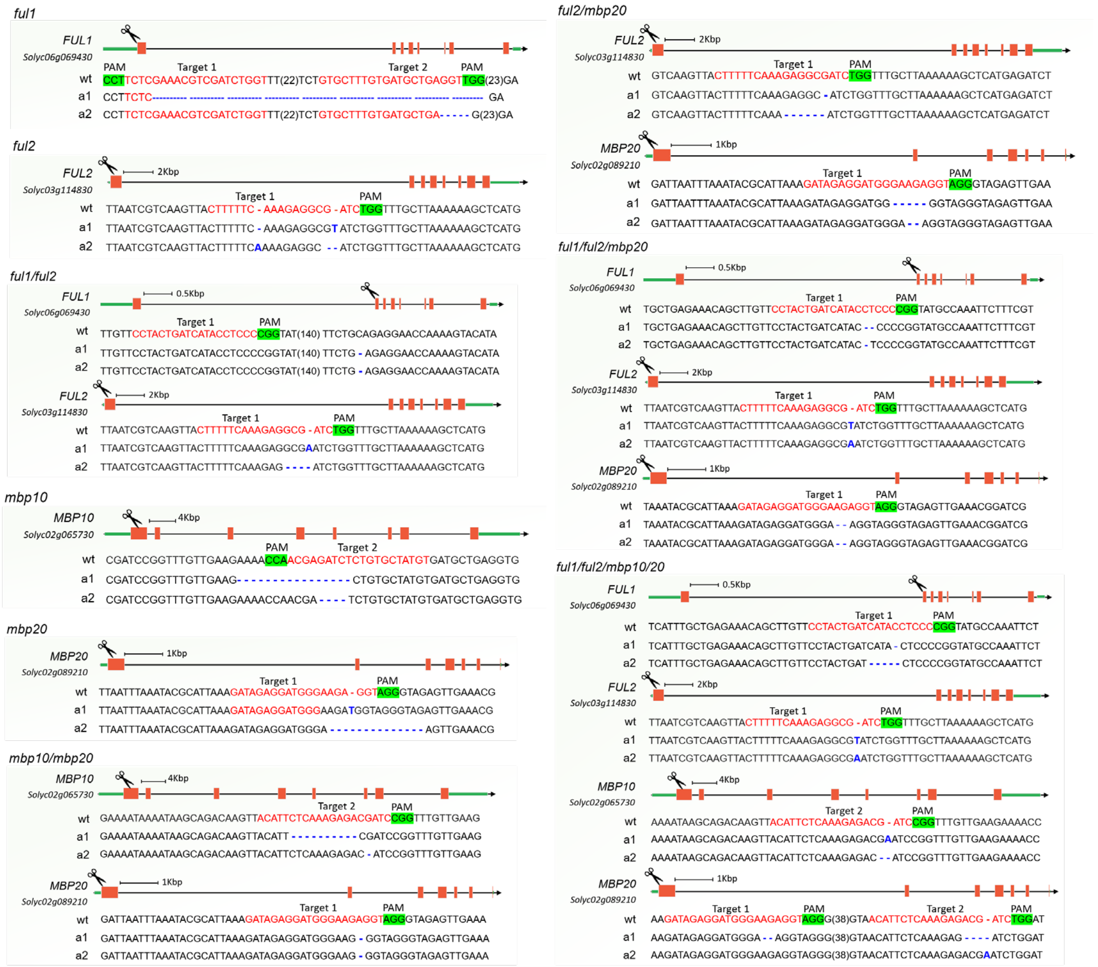

**Supplemental Figure 3**. **Mutations of tomato *FUL*-like genes generated by CRISPR/Cas9**. Sequencing of T1 transgenic progeny plants showed random insertion and deletion in target regions, resulting in two independent null loss-of-function alleles (a1 and a2) for each mutant. The orange and green boxes indicate exons and UTR of the genes, respectively. Cartoon scissor indicates the target exons. The red and green font highlights sgRNA targets and protospacer-adjacent motif (PAM) sequences, respectively. Numbers in parentheses show gap lengths. Blue dash and letter indicate deletion and insertion. The *ful2* single mutant and *ful1/ful2* double mutant lines were obtained from Wang et al., 2019.

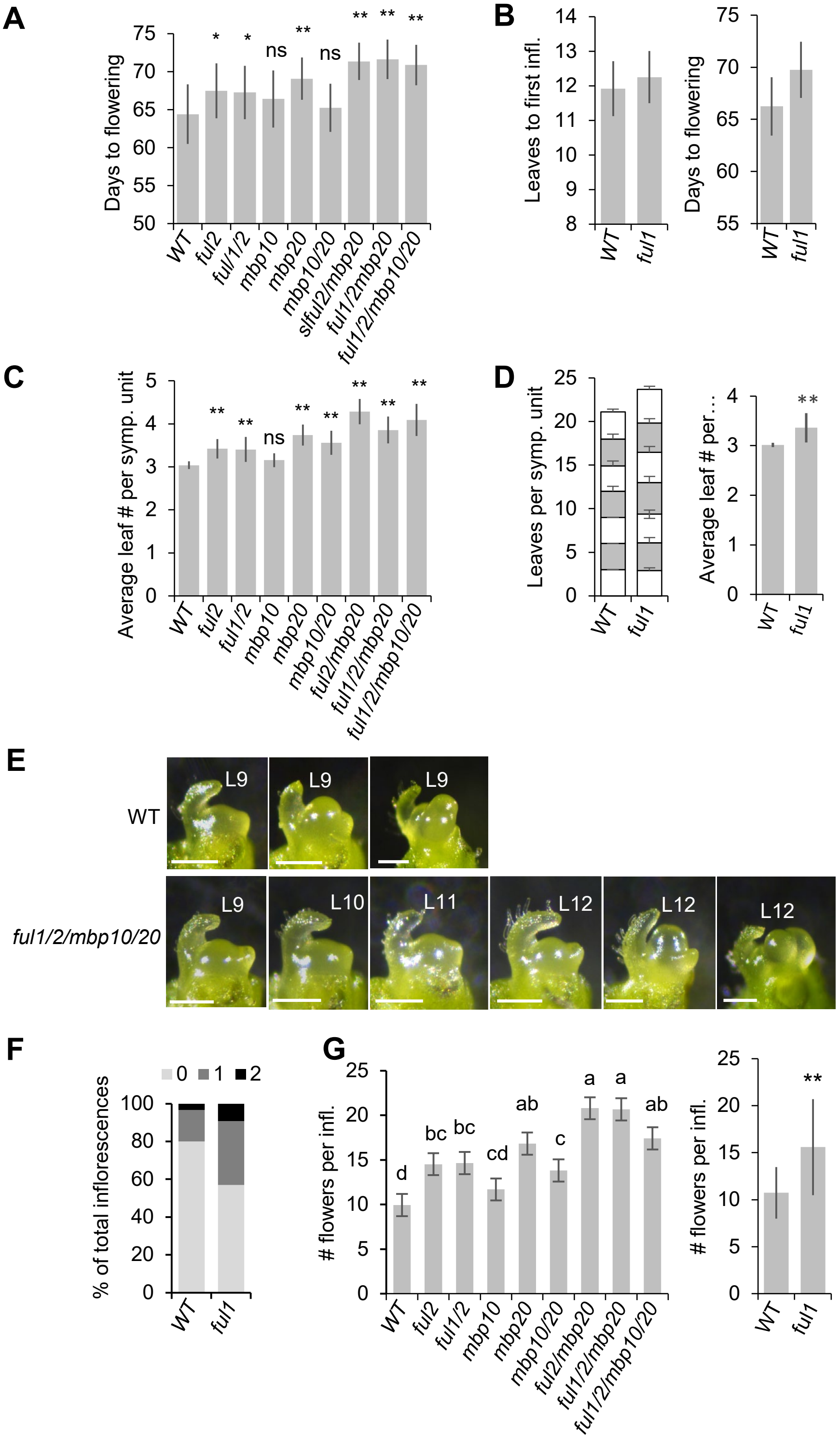
A
Days to flowering
75
70
65
60
55
50
*
*
ns
**
ns
**
**
**
WT
ful2
ful1/2
mbp10
mbp20
mbp10/20
slful2/mbp20
ful1/2mbp20
ful1/2/mbp10/20
B
Leaves to first infl.
14
13
12
11
10
9
8
WT
ful1
Days to flowering
75
70
65
60
55
WT
ful1
C
Average leaf # per symp. unit
5
4
3
2
1
0
**
**
ns
**
**
**
**
**
WT
ful2
ful1/2
mbp10
mbp20
mbp10/20
ful2/mbp20
ful1/2/mbp20
ful1/2/mbp10/20
D
Leaves per symp. unit
25
20
15
10
5
0
WT
ful1
Average leaf # per…
4
3
2
1
0
**
WT
ful1
E
WT
L9
L9
L9
ful1/2/mbp10/20
L9
L10
L11
L12
L12
L12
F
0
1
2
% of total inflorescences
100
80
60
40
20
0
WT
ful1
G
### flowers per infl.
25
20
15
10
5
0
d
bc
bc
cd
ab
c
a
a
ab
WT
ful2
ful1/2
mbp10
mbp20
mbp10/20
ful2/mbp20
ful1/2/mbp20
ful1/2/mbp10/20
### flowers per infl.
25
20
15
10
5
0
**
WT
ful1

**Supplemental Figure 4**. ***slful* mutants show delayed flowering in primary shoot and sympodial shoot**. **(A**) Quantification of primary shoot flowering times indicated by days from seed sowing to first flower opening. **(B)** Flowering data of primary shoots for wild-type (WT) and the *ful-1* single mutant. **(C)** Average leaf number of the first five successive sympodial shoots. **(D)** Flowering data of sympodial shoots for WT and the *ful-1* single mutant. **(E)** A developmental series of shoot apical meristem of WT and quadruple mutant from the vegetative stage to floral transition. White bar: 200 μm. **(F)** Inflorescence branching phenotype of WT and *ful1* mutant plants. **(G)** The quantification of the number of flowers per inflorescence. The data of B, D, F, right panel of G were observed in independent experiment. Mean values (± SD) were compared to WT using one way ANOVA followed by a post hoc LSD test. Significant differences are represented by asterisks. (∗) P-value < 0.05; (∗∗) P-value < 0.01; ns, not significant. Six individual T2 offspring plants were analyzed per line and the data from the two different genotypes were combined for each mutant (e.g. 2x6 individuals for *ful2* etc.).

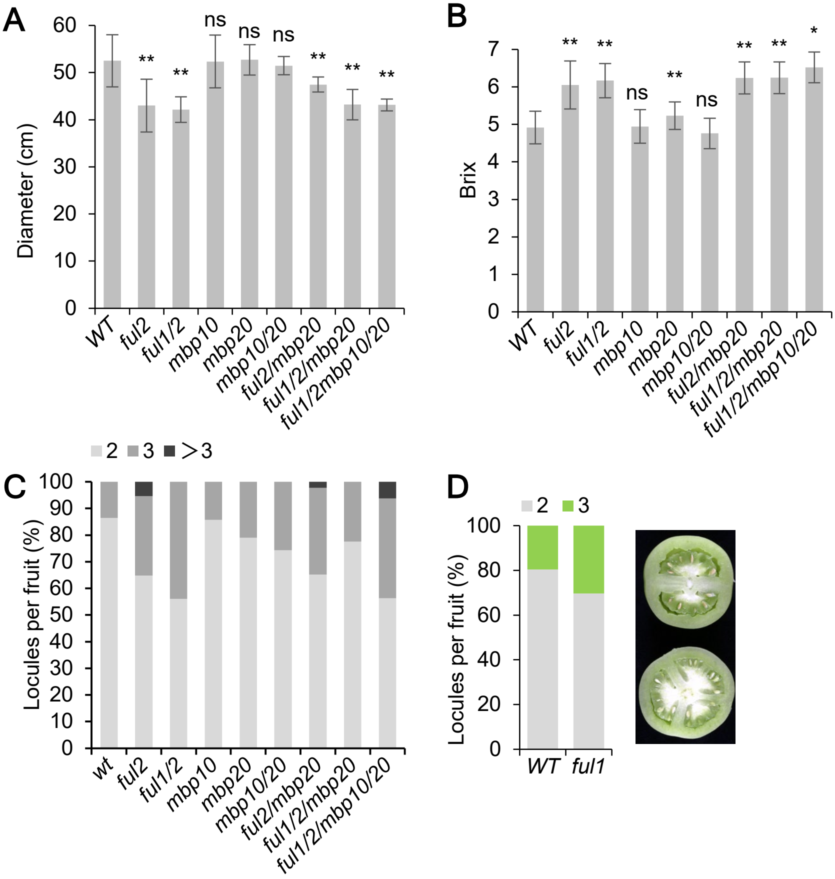

**Supplemental Figure 5. Fruit phenotypes of the *slful* mutants. (A)** Fruit diameter of wild-type (WT) and mutants. (**B**) Brix values of WT and mutants, indicating the soluble sugar contents in the fruits. Values of 16 to 40 fruits for each genotype were used. Mean values (± SD) were compared to WT using one way ANOVA. Significant differences are represented by asterisks. (*) P-value < 0.05; (**) P-value < 0.01; ns, not significant. **(C)** Proportion of fruits with a given locule number. 16-40 fruits per genotype were used.

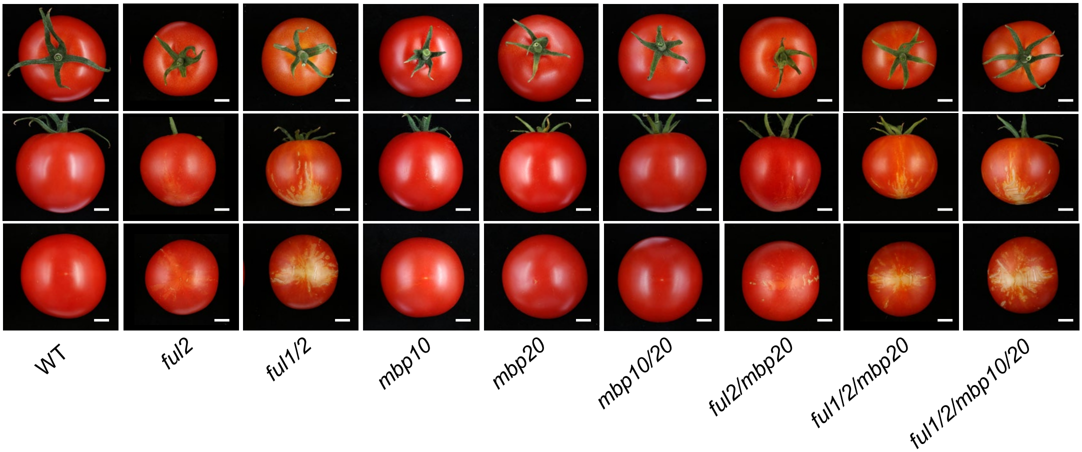

**Supplemental Figure 6**. **Fruit phenotype of wild-type and *slful* mutants**. Red ripe fruits were harvested for the picture. White bar: 1 cm.

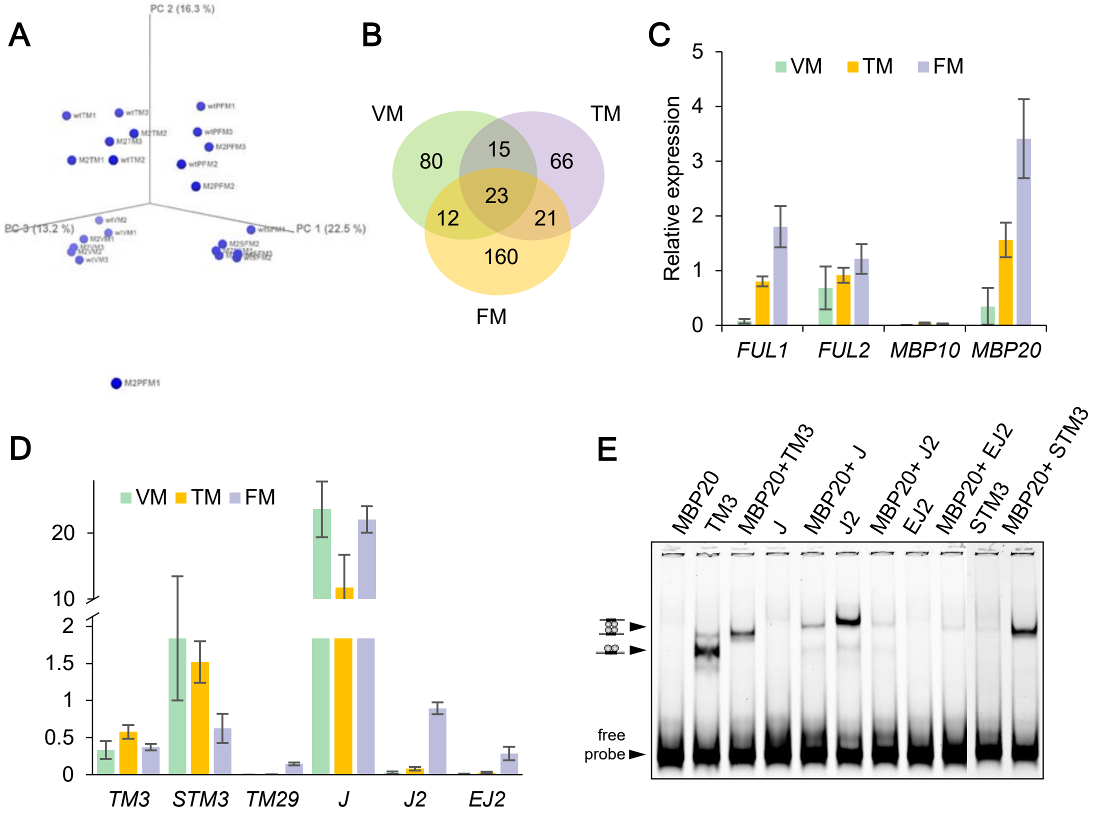

**Supplemental Figure 7. Relative expression profiles of *SlFUL* and the MADS-box genes encoding their interacting proteins. (A)** PCA of differentially expressed genes (DEGs) in the VM, TM, PFM, and SFM of WT and *ful1/2/mbp10/20* (M2), determined by RNA-seq. **(B)** Venn diagram showing the overlap of DEGs in VM, TM and FM of WT and M2**. (C), (D)** Expression of *SlFUL* genes and MADS-box genes encoding *SlFUL* interactors in three successive stages of SAM transition obtained by qRT-PCR (dCT). The values shown (mean ± SD) are the average of three replicates. VM: vegetative meristem; TM: transition meristem; FM: floral meristem.

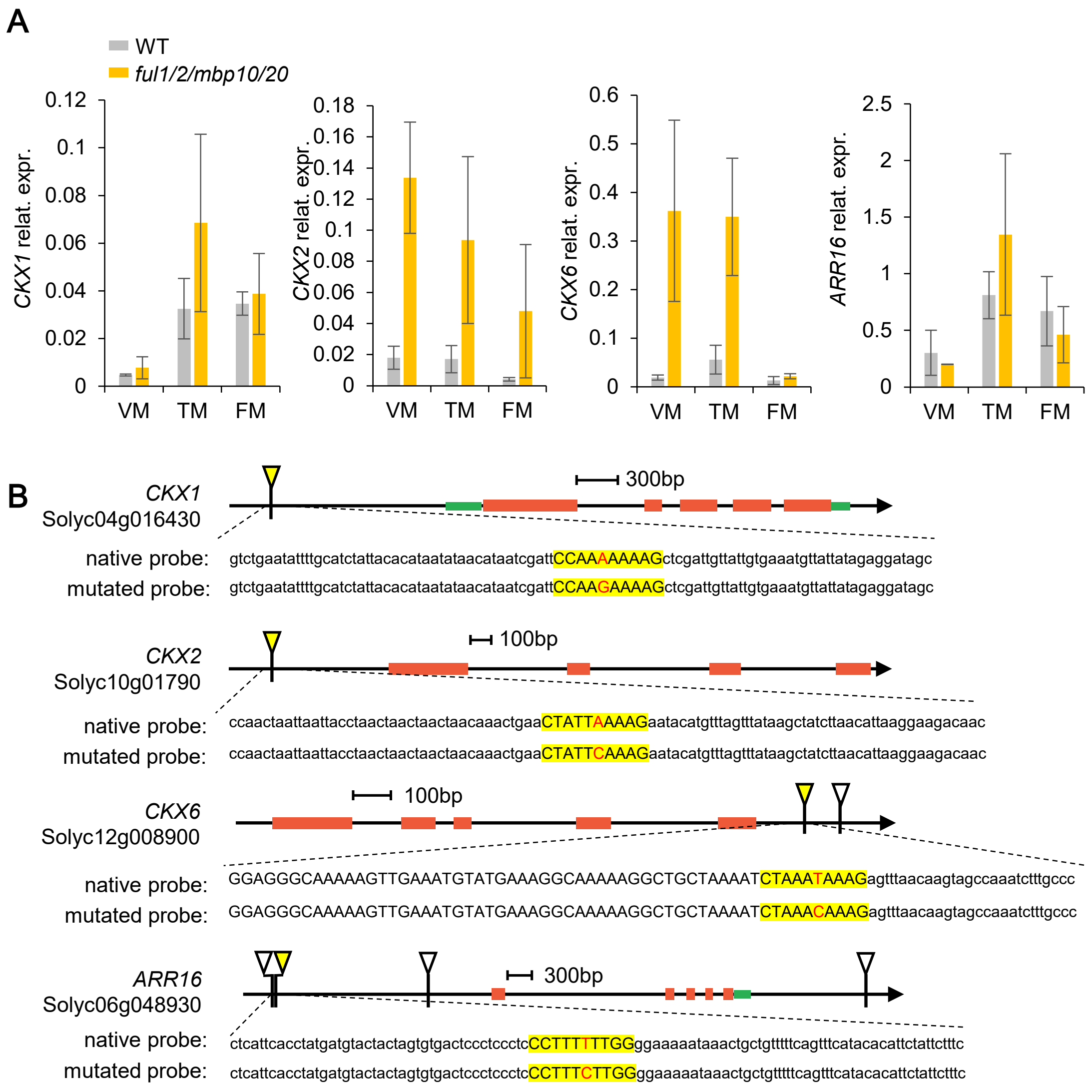

**Supplemental Figure 8. FUL2/MBP20 regulation of cytokinin negative regulators. (A)** Expression *CKX1/2/3 and ARR16* in meristems of wild-type and quadruple mutant by qPCR (dCT). VM: vegetative meristem; TM: transition meristem; FM: floral meristem. The values shown (mean ± SD) are the average of three replicates. **(B)** Schematic representation of *CKX1/2/3 and ARR16* genomic loci showing the FUL2 and MBP20 binding sequences of CArG motifs. The red and green boxes indicate exons and UTRs of genes, respectively. Triangles indicate CArG boxes and yellow-highlighted regions were tested by EMSAs. Nucleotides in red were changed in the mutated probes.

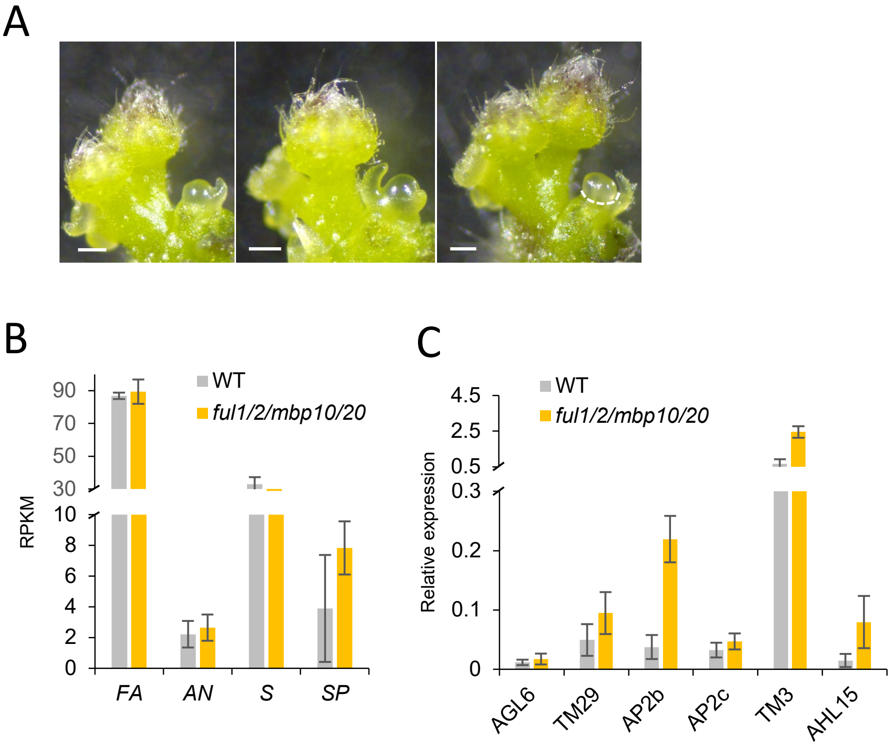

**Supplemental Figure 9. Gene expression in the floral meristem of sympodial shoot. (A)** Manual microdissection of the first floral meristem of sympodial shoot (SFM) for transcriptome profiling. Dashed lines indicate the dissected tissues. White bar: 200 µm. **(B)** Normalized gene expression (RPKM) of *FA*, *AN*, *S*, *SP* in SFM of wild-type and quadruple mutant. **(C)** Expression of AGL6, TM29, AP2b, AP2c, *TM3* and *AHL15* in the SFM of wild-type and quadruple mutant obtained by qRT-PCR **(dCT).**

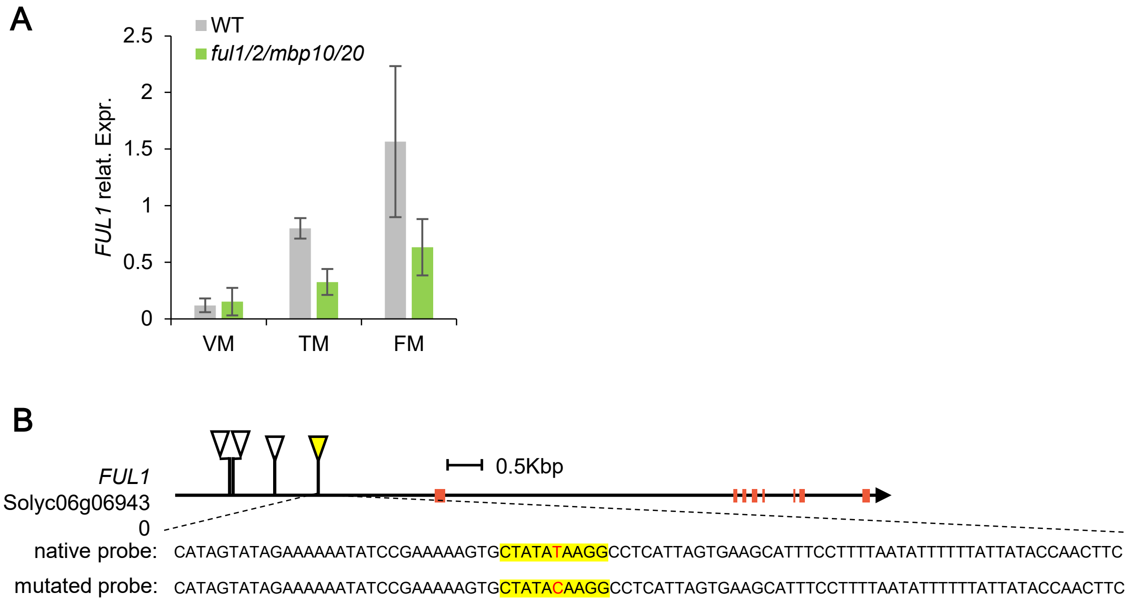

**Supplemental Figure 10. FUL2/MBP20 regulation of FUL1. (A)** *FUL1* expression in meristems of wild-type and quadruple mutant by qPCR. **(B)** Schematic representation of *FUL1* genomic loci showing the FUL2 and MBP20 binding sequences of CArG motifs. The red boxes indicate exons of genes. Triangles indicate CArG boxes and yellow-highlighted regions were tested by EMSAs. Nucleotides in red were changed in the mutated probes.
