## Supplemental Table 2 for "FRUITFULL-like genes regulate flowering time and inflorescence architecture in tomato"

Supplemental Table 2. Primer used in this study

| Gene name | Forward primer | Reverse primer |
| --- | --- | --- |
| FUL1 | AAAAAGCAGGCTTGGGAAGAGGAAGAGTCCA | AGAAAGCTGGGTTTAATTATTAAGATGACGAAGCA |
| FUL2 | AAAAAGCAGGCTTGGGTAGAGGAAGAGTACA | AGAAAGCTGGGTTTAACCGTTGAGATGTCGAAGCA |
| MBP10 | AAAAAGCAGGCTTGGGGCGGGGTAGGGTG | AGAAAGCTGGGTTCATCCTTTGTTGTGGACATGGT |
| MBP20 | AAAAAGCAGGCTTGGGAAGAGGTAGGGTAGAGT | AGAAAGCTGGGTTCATCCTTCGTTGCTGACGTGG |
| SlMBP24 | AAAAAGCAGGCTTGATGGTGAGACAAAAAATTCAG | AGAAAGCTGGGTCTAAGGAAAAGCCAAGCATAACTTG |
| J | AAAAAGCAGGCTTGGCTAGAGAAAAAATTCAGATCAAGAAAATA | AGAAAGCTGGGTTTACTTTTTTTTTTCTCCTTCTTCTAATAACATACAAAAATAG |
| TAG1 | AAAAAGCAGGCTATATGGACTTCCAAAGTGATCTAACC | AGAAAGCTGGGTTTAGACTAGTTGAATAGGGGGTTGG |
| TM5 | CACCATGGGAAGGGGTAGGGTTGAGCT | TCAAGGCAACCAGCCAGCCA |
| TM29 | CACCATGGGTAGAGGAAGAGTTGAGCTGA | TCACAGCATCCAACCAGGTATCA |
| MADS-RIN | CACCATGGGTAGAGGGAAAGTAGAATTGAAG | TCAAAGCATCCATCCAGGTACAA |
| MADS1/EJ2 | CACCATGGGAAGAGGAAGAGTTGAGCTTAAG | TTAAAGCATCCATCCATGAATAAATC |
| SlMBP21/J2 | CACCATGGGAAGAGGAAGAGTAGAACTAAAGAG | TTAGAGCATCCACCCTGGAATAAATC |
| TM3 | AAAAAGCAGGCTTGGTTCGAGGTAAAACCCAG | AGAAAGCTGGGTTAGAGACGTCTTTCTCTGCAC |
| STM3 | AAAAAGCAGGCTTGGTTCGAGGTAAAACCCAGA | AGAAAGCTGGGTAAGGGCTAAAAGTTAAACTCCATC |
| SlMBP18/FYFL | AAAAAGCAGGCTTGGTGAGAGGAAAAGTAGAAATG | AGAAAGCTGGGTCTATAAGCAGCGCATTTGAG |
| SlMBP13 | AAAAAGCAGGCTTGGTGAGAGGAAAAACTGAGT | AGAAAGCTGGGTTTAAAGTGATGTAGGTAGTGTG |
| SlMBP14 | AAAAAGCAGGCTTGGTGAGGGGAAAAACTGAG | AGAAAGCTGGGTTCATCTTTCAGGAAGTCCAA |
| SlMBP9 | AAAAAGCAGGCTTGGGGAGAGGTAAGATAGTG | AGAAAGCTGGGTGGTTTTCACTGCGAATGC |
| SlMBP12 | AAAAAGCAGGCTTGGGGAGAGGAAAGATATTG | AGAAAGCTGGGTTCCACTAATGCAGTGACAG |
| SlMBP22 | AAAAAGCAGGCTTGGGGAGAGGAAAGATAG | AGAAAGCTGGGTTCAATCGTAGGTAGAGGGAG |
| MADS-MC | AAAAAGCAGGCTTGGGAAGAGGAAAAGTTG | AGAAAGCTGGGTTCATAGATGTTTATTCATGTT |

Yeast two-hybrid assays

| Gene name | Forward primer | Reverse primer |
| --- | --- | --- |
| FUL1 | GTTTTGCCACAACAACTGGACTC | CTTGCTGCTGTGAAGAACTACC |
| FUL2 | GATAACATAATATTGTCCGCTTGC | GATAACATAATATTGTCCGCTTGC |
|  | CATGAGATCTCTGTGCTTTGCG | ATCCTTTCCATGCAAGAGTCAGT |
| MBP10 | GAATTTCGGGGTTTAGAGAAACAGC | GCTGGGAAATGGACTCGTGC |
| MBP20 | CACATTCTCACCACCAACTTCCTAA | AGTGATGAGCCTGACCCGAT |
| J | ATTGATCCTCCTCCTCAAGATGATG | CTCTTCAGCCTGAGTAAGGTAGCC |
| J2 | GCTGGCAGACCTTCAAGAAAAG | CCATTGTTTGTCCTCCATTATTTCC |
| EJ2 | AAGCAAATCAGGTCAAGGAAGACAC | CCTCCATCTTCCCAACACAATCG |
| TM3 | GGAAGTTCTTGGGAGAAGGTCTG | AGTCGCTCAACCTGTTCCTTG |
| STM3 | GCAATTGGAGCAGAGTGTCA | TCGTCTCTCTTCATCTCCTCCA |
| TM29 | GTGGAATGAGGCTAACAAGGTTCT | GACTTTGCTCACCACCACCC |
| AGL6 | TGTGAGGCTGAAGTTGCTCT | GCAACAACGTTGGTACCTCTC |
| AP2b | ACCCAAGCAACCTAGTCCAC | CCGGAGAATGTAGGTGCGTT |
| AP2c | TGGATATTGATTGGCAGCGC | TGGAGAATGCAAGTGCGTCT |
| AHL15 | TTGAAGTCATCCGCCGTTCA | TGGGTTTTTCCACGTGACCA |
| CKX1 | TGGGCAAGCATTCAAACATGG | AGAACAGGTCAGCATTCTGC |
| CKX2 | CTTCCAAAGATTTTGGAAAGATC | GATAGAAAGGCCATAAGAAAATTGA |
| CKX6 | CTAATGCTGGAATTAGTGGTCAAAC | CATATCTTTGGAGCAAGTCATTAAT |
| ARR16 | AAGGGCGTTGGAGTACTTGG | CCAGTCATTCCTGGCATGCA |
| CAC | CCTCCGTTGTGATGTAACTGG | ATTGGTGGAAAGTAACATCATCG |

RT-PCR

CRISPR/Cas9 genome-editing

| sgRNA name | Forward primer | Reverse primer |
| --- | --- | --- |
| FUL1-sgRNA-1 | TGTGGTCTCAATTGTGTTCTCTATTCGCTTCAACGTTTTAGAGCTAGAAATAGCAAG | TGTGGTCTCAAGCGTAATGCCAACTTTGTAC |
| FUL1-sgRNA-2 | TGTGGTCTCAATTGACCAGATCGACGTTTCGAGAGTTTTAGAGCTAGAAATAGCAAG |  |
| FUL1-sgRNA-3 | TGTGGTCTCAATTGTGCTTTGTGATGCTGAGGTGTTTTAGAGCTAGAAATAGCAAG |  |
| FUL1-sgRNA-4 | TGTGGTCTCAATTGCCTACTGATCATACCTCCCGTTTTAGAGCTAGAAATAGCAAG |  |
| FUL2-sgRNA | TGTGGTCTCAATTGCTTTTTCAAAGAGGCGATCGTTTTAGAGCTAGAAATAGCAAG |  |
| MBP10-sgRNA-1 | TGTGGTCTCAATTGAAAGATGGGGCGGGGTAGGGGTTTTAGAGCTAGAAATAGCAAG |  |
| MBP10-sgRNA-2 | TGTGGTCTCAATTGACATAGCACAGAGATCTCGTGTTTTAGAGCTAGAAATAGCAAG |  |
| MBP10-sgRNA-3 | TGTGGTCTCAATTGATATGAAAATTACTCATACGGTTTTAGAGCTAGAAATAGCAAG |  |
| MBP20-sgRNA-1 | TGTGGTCTCAATTGATAGAGGATGGGAAGAGGTGTTTTAGAGCTAGAAATAGCAAG |  |
| MBP20-sgRNA-2 | TGTGGTCTCAATTGTTGATTGTGTTTTCTACCAAGTTTTAGAGCTAGAAATAGCAAG |  |
| MBP20-sgRNA-3 | TGTGGTCTCAATTGACATTCTCAAAGAGACGATCGTTTTAGAGCTAGAAATAGCAAG |  |
| MBP10/20-sgRNA | TGTGGTCTCAATTGACATTCTCAAAGAGACGATCGTTTTAGAGCTAGAAATAGCAAG |  |

| Gene name | Forward primer | Reverse primer |
| --- | --- | --- |
| FUL1 | GACCTTCGCTTATAGCTCTATCCC | CTTCTCCCACATAATGCCTGC |
| FUL2 | CTACCTGGGGAGATCCTTCC | TGAGTCCAACTTCAGCATCG |
| MBP10 | TGTTGCTCTCCTGCATAGCA | ACAAATTGAAGATGGAGAACGGAG |
| MBP20 | CCAATCACAAATCGACAACGCAAC | ATGAGGACATCAATGAGCTGATC |
| Cas9 | CTTTGGCAATATCGTGGACG | CGTTCTTCTTCTCCCCAGGG |
| NPT2 | AGACAATCGGCTGCTCTGAT | AGCCAACGCTATGTCCTGAT |

Genotyping PCR

| Gene | Forward primer | Reverse primer | Description |
| --- | --- | --- | --- |
| FUL1-dC | AAAAAGCAGGCTATGGGAAGAGGAAGAGTCCAG | AGAAAGCTGGGTTTACATTAGTACTCTGGTATG | BP reaction |
| FUL2 | AAAAAGCAGGCTATGGGTAGAGGAAGAGTACA | AGAAAGCTGGGTTTAACCGTTGAGATGTCGAAGCA | BP reaction |
| MBP20 | AAAAAGCAGGCTATGGGAAGAGGTAGGGTAG | AGAAAGCTGGGTTCATCCTTCGTTGCTGACGTGG | BP reaction |
| TM3 | AAAAAGCAGGCTATGGTTCGAGGTAAAACCCAGA | AGAAAGCTGGGTTAGAGACGTCTTTCTCTGCAC | BP reaction |
| STM3 | AAAAAGCAGGCTATGGTTCGAGGTAAAACCCAG | AGAAAGCTGGGTTCAAGACCATTCAGGACGCC | BP reaction |
| J | AAAAAGCAGGCTATGGCTAGAGAAAAAATTCAGATC | AGAAAGCTGGGTTTACTTTTTTTTTTCTCCTTCTTCTAATAACATACAAAAATAG | BP reaction |
| J2 | AAAAAGCAGGCTATGGGAAGAGGAAGAGTAGAAC | AGAAAGCTGGGTTTAGAGCATCCACCCTGGAA | BP reaction |
| EJ2 | AAAAAGCAGGCTATGGGAAGAGGAAGAGTTGAGC | AGAAAGCTGGGTTTAAAGCATCCATCCATGAA | BP reaction |
| ARR16 | CTCATTCACCTATGATGTAC | GAAAGAATAGAATGTGTATGAAACTG | Probe overlap PCR |
| mARR16 | TCCCTCCCTCCCTTTCTTGGGG | CCCCAAGAAAGGGAGGGAGGGA |  |
| CKX1 | GTCTGAATATTTTGCATCTATTAC | GCTATCCTCTATAATAACATTTCAC | Probe overlap PCR |
| mCKX1 | CGATTCCAAGAAAAGCTCG | CGAGCTTTTCTTGGAATCG |  |
| CKX2 | CCAACTAATTAATTACCTAACTAACTAAC | GTTGTCTTCCTTAATGTTAAGATA | Probe overlap PCR |
| mCKX2 | CTGAACTATTCAAAGAATAC | GTATTCTTTGAATAGTTCAG |  |
| CKX6 | GGAGGGCAAAAAGTTGAAATG | GGGCAAAGATTTGGCTACTT | Probe overlap PCR |
| mCKX6 | GGCTGCTAAAATCTAAACAAAG | CTTTGTTTAGATTTTAGCAGCC |  |
| FUL1 | CATAGTATAGAAAAAATATCCG | GAAGTTGGTATAATAAAAAATATTAAAAG | Probe overlap PCR |
| mFUL1 | CTAATGAGGCCTTGTATAGCAC | GTGCTATACAAGGCCTCATTAG |  |

EMSAs
